# Light-harvesting by antenna-containing xanthorhodopsin from an Antarctic cyanobacterium

**DOI:** 10.1101/2025.06.10.658827

**Authors:** María del Carmen Marín, Shunya Murakoshi, Andrey Rozenberg, Tatsuki Tanaka, Masae Konno, Wataru Shihoya, Osamu Nureki, Keiichi Inoue, Oded Béjà

## Abstract

Microbial rhodopsins are light-sensitive proteins vital to various phototrophic and sensory processes in microorganisms. Xanthorhodopsins, with their dual chromophore system involving retinal and carotenoids, have been predominantly studied in the extreme halophilic bacterium *Salinibacter ruber* and in the primitive thylakoid-less cyanobacterium *Gloeobacter violaceus*, where they facilitate light-driven outward proton pumping and enhanced light-harvesting. However, their distribution, binding specificity, and ecological significance in cyanobacteria remain poorly understood. Here we report widespread xanthorhodopsin genes in cyanobacterial genomes and characterize a protein from an Antarctic filamentous cyanobacterium that uniquely binds hydroxylated lutein. Through bioinformatics, spectroscopic, functional and structural assays analyses, we determine the properties and ecological role of this cyanobacterial xanthorhodopsin. Our findings highlight xanthorhodopsins’ role in modulating light-harvesting efficiency in cyanobacteria, particularly in extreme environments. The lutein binding and associated structural changes likely provide selective advantages for adapting to polar light conditions, contributing to cyanobacterial survival in harsh habitats.

## Introduction

Microbial rhodopsins, a superfamily of heptahelical photoreceptive membrane proteins using a retinal chromophore, are crucial to understanding how microorganisms adapt to and harness light energy. These proteins exhibit a remarkable structural and functional diversity^1–3^, which enables their application in optogenetics and synthetic biology. Among these, some members of the proteorhodopsin (PR) and xanthorhodopsin (XR) families stand out due to their unique dual chromophore system, which includes a retinal moiety and a non-covalent carotenoid antenna. So far, since the discovery of *Sr*XR, the first antenna-binding XR from the extreme halophilic bacterium *Salinibacter ruber*, only a few bacterial PRs and XRs have been demonstrated to form such complexes^4–8^. One of the best studied proton-pumping rhodopsins with a carotenoid antenna is GR from the thylakoid-less terrestrial cyanobacterium *Gloeobacter violaceus*^6^ which, similarly to *Sr*XR, recruits a 4-ketocarotenoid antenna (echinenone) to transfer light energy to the retinal chromophore^9^. Nevertheless, increasing evidence suggests that in contrast to the classical ketocarotenoid-binding rhodopsins GR and *Sr*XR, rhodopsin proton pumps binding hydroxylated carotenoids (xanthophylls) might be more common in aquatic environments^7,8,10^.

The appearance of ion-pumping rhodopsins in organisms with chlorophyll-based photosystems is well documented, although their physiological roles and the interplay between the two types of photosystems are known only in isolated cases^7,11–15^. Recently, a possible connection between anoxygenic photosynthesis and rhodopsin-based phototrophy was suggested by Kopejtka and coworkers^7^, who described a freshwater alphaproteobaterium capable of using two different systems for light harvesting: a proton pumping rhodopsin and an anoxygenic photosynthetic apparatus with bacteriochlorophyll in the reaction center. Interestingly, both systems were shown to use the same carotenoid (nostoxanthin) as an antenna, highlighting a potential evolutionary strategy for optimizing light capture in diverse phototrophic processes. This overlap in antenna systems suggests a functional convergence where organisms leverage similar molecular components across distinct photosystems, potentially facilitating efficient energy transfer and flexibility in response to varying light conditions.

Cyanobacteria play a crucial role in aquatic and terrestrial ecosystems through oxygenic photosynthesis, contributing significantly to primary production and biogeochemical cycles^16^. The current study aims to elucidate the distribution of antenna- possessing rhodopsins among cyanobacteria beyond GR. We report the discovery of a novel XR in an Antarctic cyanobacterium (cyXR) that binds a hydroxylated carotenoid. Through bioinformatic analysis, spectroscopic techniques, and functional assays, we have characterized cyXR and its potential role in enhancing cyanobacterial adaptation to diverse light environments. Our findings suggest that this novel XR compliments oxygenic photosynthesis (separately or in parallel) in Antarctic cyanobacteria, thereby contributing to the ecological success of cyanobacteria.

## Methods

### Search for rhodopsin genes in cyanobacteria

The incidence of different rhodopsin genes in cyanobacteria was estimated based on genomes in JGI’s Genomic catalog of Earth’s Microbiomes (GEMs)^17^ (genomes and operational taxonomic unit [OTU] representatives) and assemblies of GTDB r. 214^18^ species representatives. Assemblies assigned to class “*Cyanobacteriia*” were translated using getorf from EMBOSS v. 6.6.0^19^ (minimum ORF size 150 nt) and rhodopsins were identified using Pfam profile PF01036.24 with hmmsearch from HMMER v. 3.4^20^ with the default gathering cutoff. The matching proteins were classified to family using a curated database of prokaryotic regular microbial rhodopsins with blastp from NCBI BLAST v. 2.16.0^21^. Best matches with an E-value of <1e-10 and ≥50% identity to the reference were used for family assignment. The results were visualized using ComplexUpset v. 1.3.3^22^.

### Phylogenetic analysis

For the phylogeny of xanthorhodopsins across bacterial taxa the following strategy was utilized. *Sr*XR was used to search JGI GEM genomes and OTU representatives and assemblies of GTDB species representative with tblastn from NCBI BLAST. Genes on matching scaffolds (E-value threshold of 1e-10 and at least 200 residues alignment length) were predicted with prodigal v. 2.6.3^23^ in metagenomic mode. To extract XR sequences, the resulting proteins were searched against the curated rhodopsin database (see above) with blastp and best matches to reference XRs with an E-value threshold of 1e-30 and ≥40% identity were retained. The XRs were combined with outgroup sequences (NQ pumps and clade P1 representatives *Qs*ActR, *Kr*ActR, and DSE009) and reference XRs, clustered at 95% identity level with CD-HIT v. 4.8.1^24^, aligned with MAFFT v. 7.525^25^ in automatic mode, trimmed with trimAl v. 1.4.1^26^ allowing up to 10% gaps per position and phylogeny was reconstructed with IQ-TREE v. 2.2.2.3^27^ with 1000 replicates for ultra-fast bootstrap^28^. In the resulting phylogeny, the ingroup appeared monophyletic (confirming the classification of the sequences) and the outgroups were used for rooting. For the fine-grained phylogenetic analysis, sequences belonging to the cyanobacterial XR clade were extracted from a database containing rhodopsins from JGI GEM genomes, GTDB species representatives, proteins from JGI’s Integrated Microbial Genomes & Microbiomes (IMG/M)^29^ assigned to the Pfam family PF01036 (downloaded in November 2023) and the UniRef100 r. 2023_05^30^ database based on similarity to the initially collected cyanobacterial XRs and preliminary phylogenetic analyses. Sequences at least 220 residues in length were combined with outgroup XRs, clustered at 100% identity level with CD-HIT, aligned with MAFFT in automatic mode, trimmed with trimAI in atomated1 mode and the phylogeny was reconstructed using RAxML v. 8.2.12^31^ under the PROTGAMMAAUTO model and with 1000 rapid bootstrap replicates. The corresponding cyanobacterial assemblies were taxonomically classified using GTDBtk v. 2.3.2^32^ with GTDB r. 214 as a reference database.

### XR gene neighbors

Regions of at most 20,000 nt surrounding the XR genes were extracted and annotated using PGAP v. 2024-04-27.build7426^33^. Immediate neighbors chosen for further annotation included genes and operons within ca. 1800 nt of the XR gene. Representative genomic fragments were chosen per cluster of similar genomic contexts and homologous genes were identified with Proteinortho v. 6.0.25^34^ and based on the functional annotations.

### Construction of DNA plasmids expression

The gene chosen for expression originated from JGI GEM genome 3300012044_11, an Antarctic MAG from Dry Valleys (JGI Genomes Online Database analysis Ga0136636) classified in the genus *Pseudanabaena* (*Pseudanabaenales*: *Pseudanabaenaceae*). cyXR was codon-optimized for expression in *Escherichia coli*, synthesized (GenScript, China), and cloned into pET21a(+) vector (Novagen, Merck KGaA, Germany) with a C-terminal 6×His-tag, using NdeI and XhoI restriction sites.

### Protein expression and purification

*E. coli* cells harboring the cyXR-cloned plasmid were cultured in 2×YT medium containing 50 μg/mL ampicillin. The expression of C-terminal 6× His-tagged protein was induced by 0.1 mM isopropyl-𝛽-D-thiogalactopyranoside (IPTG) in the presence of 10 μM all-*trans*- retinal (Toronto Research Chemicals, Canada) at 37 °C for 4 h. The harvested cells were sonicated (Ultrasonic Homogenizer VP-300N; TAITEC, Japan) for disruption in a buffer containing 50 mM Tris–HCl (pH 8.0) and 5 mM MgCl_2_. The membrane fraction was collected through ultracentrifugation (CP80NX; Eppendorf Himac Technologies, Japan) at 142,000 ×*g* for 1 h. The protein was solubilized in a buffer containing 50 mM MES– NaOH (pH 6.5), 300 mM NaCl, 5 mM imidazole, 5 mM MgCl_2_, and 3% *n*-dodecyl-𝛽-D- maltopyranoside (DDM) (ULTROL Grade; Calbiochem, Sigma-Aldrich, MO). Solubilized protein was separated from an insoluble fraction through ultracentrifugation at 142,000 ×*g* for 1 h. Protein was purified using a Co-NTA affinity column (HiTrap TALON crude; Cytiva, MA). The column was washed with a 15-column volume buffer containing 50 mM MES–NaOH (pH 6.5), 300 mM NaCl, 50 mM imidazole, 5 mM MgCl_2_, and 0.1% DDM. Protein was eluted in a buffer containing 50 mM Tris–HCl (pH 7.0), 300 mM NaCl, 300 mM imidazole, 5 mM MgCl_2_, and 0.1% DDM. Eluteined protein was immediately concentrated using a 50-mL centrifugal ultrafiltration filter with a 30-kDa molecular weight cutoff (Amicon Ultra-4, Millipore, Merck KGaA, Germany), and the sample was dialyzed against a buffer containing 50 mM HEPES–NaOH (pH 7.0), 150 mM NaCl, 10% glycerol, and 0.1% DDM.

### Proton transport activity assay

Rhodopsin-expressing *E. coli* cells were collected through centrifugation at 4,800 ×*g* at 20 °C for 2 min (CF15RF; Eppendorf Himac Technologies, Japan) and washed with 100 mM NaCl. The cells were equilibrated thrice with rotational mixing in 100 mM NaCl at room temperature for 10 min. Finally, the cells were suspended in 7.5 mL of unbuffered 100 mM NaCl, and the optical density (OD) at 600 nm was adjusted to 2. The cell suspension was placed and stirred in the dark in a glass cell at 5, 12.5, and 20 °C and illuminated at *λ* = 550 ± 5 nm from the output of a 300 W xenon light source (MAX-303, Asahi Spectra, Japan) through a bandpass filter (HMX0550, Asahi Spectra, Japan). Light-induced pH changes were measured using a pH electrode (9618S-10D, HORIBA, Japan). The measurements were repeated under the same conditions after the addition of carbonyl cyanide *m*-chlorophenylhydrazone (CCCP, final concentration = 10 μM).

To quantitatively compare ion transport activity, the amount of protein was determined by measuring near-UV absorption of retinal oxime produced by the hydrolysis reaction between the retinal Schiff base (RSB) in the protein and hydroxylamine (HA). Briefly, rhodopsin-expressing *E. coli* cells were washed with a buffer containing 133 mM NaCl and 66.5 mM Na_2_HPO_4_ (pH 7.0). The washed cells were treated with 1 mM lysozyme and a small amount of Dnase I for 1 h and disrupted through sonication. To solubilize rhodopsins, 3% DDM was added, and the samples were stirred overnight at 4 °C. The rhodopsins were bleached with 500 mM HA and illuminated with visible light (*λ* > 500 nm) from the output of a 300 W xenon lamp (MAX-303, Asahi Spectra, Japan) through long-pass (Y-52, AGC Techno Glass, Japan) and heat-absorbing (HAF-50S-50H, SIGMAKOKI, Japan) filters. The absorption changes due to the bleaching of rhodopsin by the hydrolysis reaction between retinal and HA and the formation of retinal oxime were measured using a UV-visible spectrometer (V-750, JASCO, Japan). The amount of rhodopsin expressed in *E. coli* cells was estimated by the absorbance of produced retinal oxime and its differential molar extinction coefficient (*ε*) (33,600 M^−1^ cm^−1^)^35^ relative to the original absorption of unhydrolyzed retinal (Fig. 2C).

**Fig. 1.**
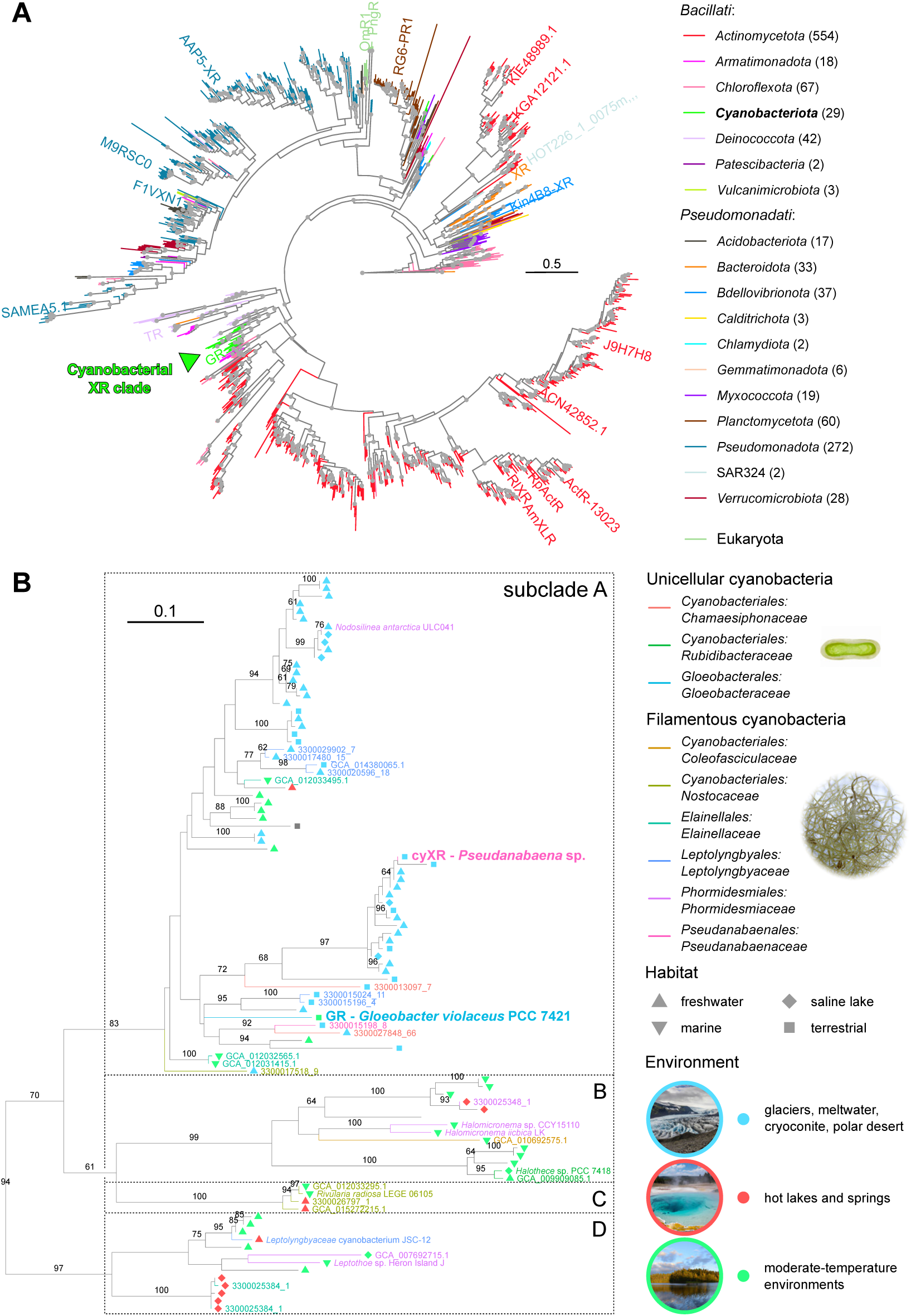
Phylogenetic relationships between xanthorhodopsins found in cyanobacteria. **A** Phylogenetic tree of XR proteins from diverse genomes classified to phylum. Proteins characterized previously or otherwise mentioned in literature are shown for reference. Numbers in parentheses next to the names of the phyla indicate the number of the representative sequences from the corresponding phylum used for the tree reconstruction. Only phyla with at least two sequences are shown. The position of the XR from cyanobacteria (phylum *Cyanobacteriota*, class “*Cyanobacteriia*”) is indicated with a green triangle. Other unrelated *Cyanobacteriota* XRs belong to the non-photosynthetic “*Ca.* Sericytochromatia”. The tree is outgroup-rooted and the outgroups are not shown. Gray dots indicate branches with ultra-fast bootstrap support values ≥95. **B** Phylogenetic relationships among the cyanobacterial XRs. Symbol shapes and colors indicate the habitats from which the corresponding isolates or MAGs originate. Labeled are sequences coming from genome assemblies: MAGs are referred to by their accession numbers and isolates are referred to by the names of the corresponding organisms. The genome labels are colored by the inferred taxonomy based on GTDBtk. Note that the inference about the morphotypes for MAGs is based on the knowledge available for isolates from the corresponding families and thus some of the MAGs from predominantly filamentous families might in reality come from secondarily unicellular cyanobacteria. The tree is outgroup-rooted and the outgroups are not shown.

**Fig. 2.**
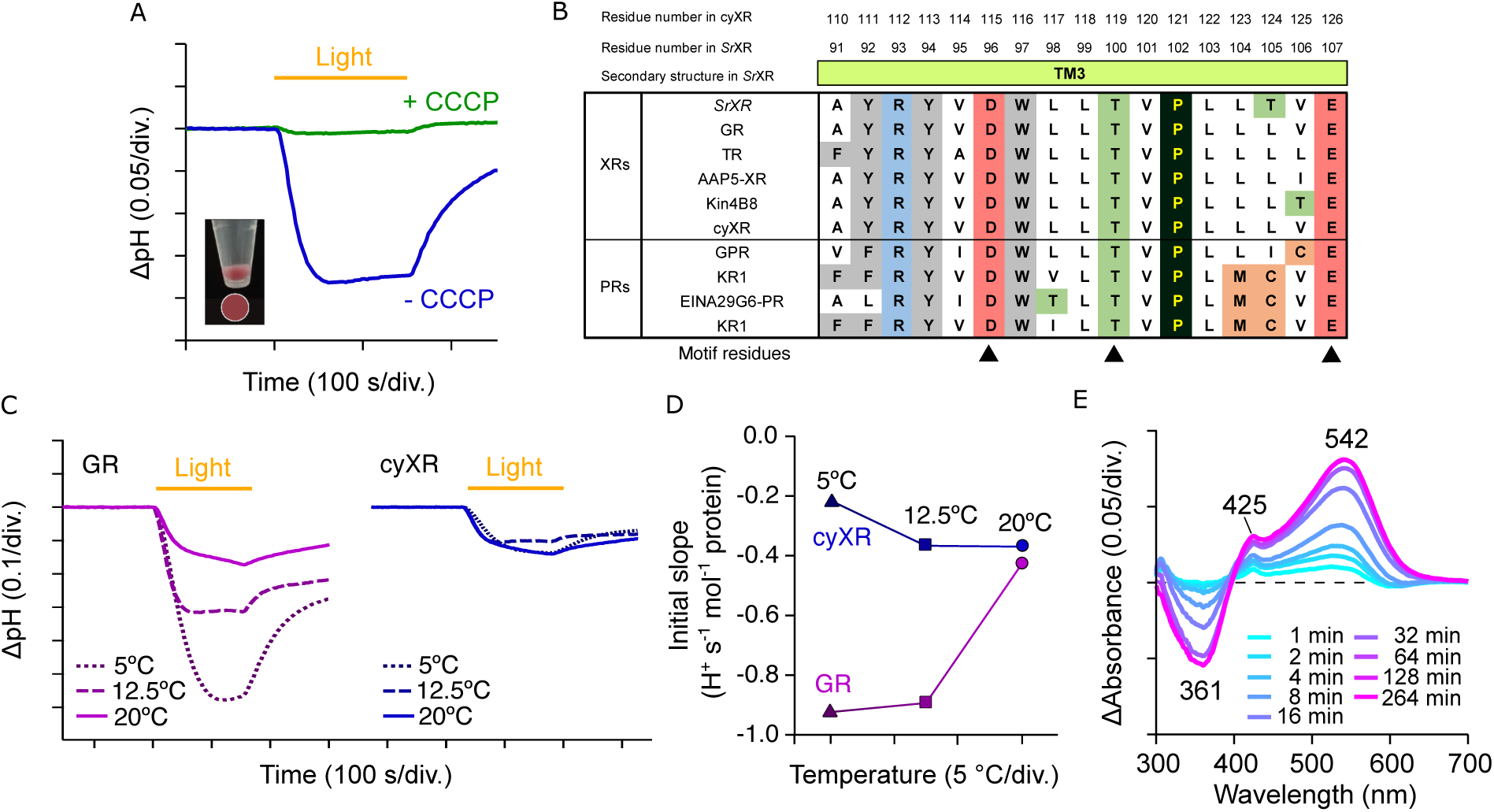
**Amino acid sequence and light-driven active proton transport of cyXR**. **A** Proton transport activity assay of cyXR, derived from an Antarctic cyanobacterium, in *E. coli* cells without (blue line) and with (green line) 10 μM CCCP. The cells were illuminated with light (*λ* > 500 nm) for 150 s (yellow line). The picture of the pellet of *E. coli* cells expressing cyXR, highlighting the strong purple color by the circle, is shown next to the result; **B** Amino acid sequence of cyXR in TM3 compared with other XRs and PRs. Black triangles indicate the motif residues; **C** Proton transport activity assay of GR (violet) and cyXR (blue) at different temperatures; **D** Initial H^+^ transport rates of GR and cyXR at different temperatures; **E** Difference in absorption spectra before and after HA bleaching reaction of cyXR in solubilized *E. coli* membranes at pH 7.0. The *λ*_max_ value was determined based on the position of the absorption peak indicated and the absorption of retinal oxime produced by the hydrolysis reaction of RSB and HA was observed as a peak at 360–370 nm.

### Purification of carotenoid-binding cyXR

Carotenoid-binding cyXR was obtained by mixing purified protein with carotenoids (lutein (Sigma-Aldrich, MO, USA) or canthaxanthin (Sigma-Aldrich, MO, USA)). First, carotenoids were dissolved in dimethyl sulfoxide (DMSO) at twice the molar amount of cyXR to be bound. The molar amounts of each were estimated by measuring the absorption spectrum (Fig. 4A), using a molar extinction coefficient of *ε* = ∼50,000 M^−1^ cm^−1^ at the absorption maximum of rhodopsin and *ε* = 145,000 M^−1^ cm^−1^ at 445 nm for carotenoids. The volume of DMSO was adjusted so that it did not exceed 2% of the rhodopsin solution volume. After preparing the carotenoid solution as described above, it was added to cyXR and left on ice and in the dark for 2 h, gently mixing the solution every 15 min. To remove the excess of carotenoids not bound to cyXR, cyXR complexed with carotenoids was purified again using the same method as the purification of the original cyXR protein. The purified protein was immediately concentrated using a 50-mL centrifugal ultrafiltration filter with a 30-kDa molecular weight cutoff (Amicon Ultra-4, Millipore, Merck KGaA, Germany), and the sample was dialyzed against a buffer containing 20 mM HEPES–NaOH (pH 7.0), 100 mM NaCl, and 0.05% DDM.

### High-performance liquid chromatography (HPLC) analysis of retinal isomers

Retinal configuration in cyXR was analyzed by HPLC using purified proteins in a buffer containing 20 mM Tris–HCl (pH 8.0), 100 mM NaCl, and 0.05% DDM. Before the measurements, the OD of the samples at their *λ*_max_ was adjusted to ∼0.2 (protein concentration of approximately 0.1 mg mL^−1^), and the proteins were stored at 4 °C overnight for dark adaptation. The HPLC system was equipped with a silica column particle size of 3 μm, 150 × 6.0 mm; Pack SIL, YMC, Japan), a pump (PU-4580, JASCO, Japan), and a UV-vis detector (UV-4570, JASCO, Japan). The solvent was composed of 15% (v/v) ethyl acetate and 0.15% (v/v) ethanol in *n*-hexane and with a flow rate of 1.0 mL min^−1^. To denature the protein, 280 μL of 90% methanol solution was added to the 75 μL sample. Retinal oxime formed by the hydrolysis reaction with 25 μL of 2 M HA solution was extracted with 800 μL of *n*-hexane, and 200 μL of the solution was injected into the HPLC system. For measurements during light illumination, the sample solutions were illuminated at *λ* = 550 ± 10 nm using a bandpass filter (AGC Techno Glass, Japan) for 1 min, followed by denaturation and hydrolysis of the retinal chromophore under illumination. For measurements of light-adapted samples, the sample solutions were illuminated at *λ* = 550 ± 10 nm using a bandpass filter (AGC Techno Glass, Japan) for 1 min, and after waiting for 1 min, denaturation and hydrolysis reactions of the retinal chromophore were performed. The molar compositions of the retinal isomers were calculated from the areas of the corresponding peaks in the HPLC patterns using the molar extinction coefficients at 360 nm for each isomer (all-*trans-*15-*syn:* 54,900 M^−1^ cm^−1^; all-*trans*-15-*anti*: 51,600 M^−1^ cm^−1^; 13--*cis-*15-*syn*: 49,000 M^−1^ cm^−1^; 13-*cis*-15*–anti*: 52,100 M^−1^ cm^−1^; 11-*cis*-15-*syn*: 35,000 M^−1^ cm^−1^; 11-*cis*-15-*anti*: 29,600 M^−1^ cm^−1^). Three independent measurements were performed to estimate experimental errors. The compositions of the retinal isomers are listed in Supplementary Table S1. Peaks were assigned by comparing the elution time with those of authentic retinal oxime isomers.

### pH titration

To investigate the pH dependence of the absorption spectra, the concentration of proteins was adjusted to OD = ∼0.5 (protein concentration of approximately 0.25 mg mL^−1^) at *λ*_max_ and solubilized in a 6-mix buffer (trisodium citrate, MES, HEPES, MOPS, CHES, CAPS (10 mM each, pH 7.0), 100 mM NaCl, and 0.05% DDM). The pH was adjusted to desired values by adding small aliquots of HCl and NaOH. Absorption spectra were recorded using a UV-vis spectrometer (V-750, JASCO, Japan). The measurements were performed at every 0.3–0.6 pH value.

### Laser-flash photolysis

For the laser-flash photolysis spectroscopy, the protein was solubilized in 20 mM HEPES–NaCl (pH 7.0), 100 mM NaCl, 0.05% DDM. OD of the rhodopsin was adjusted to approximately ∼0.4–0.5 (protein concentration of approximately 0.2–0.25 mg mL^−1^ at the *λ*_max_). The laser-flash photolysis measurement was conducted as previously described^36,37^. Nano-second laser pulses from an optical parametric oscillator (*λ*_exc_ = 550 nm, 4.5 mJ pulse^−1^ cm^−2^, 3.3 Hz (basiScan, Spectra-Physics, CA) pumped by the third harmonics of Nd-YAG laser (*λ* = 355 nm, INDI40, Spectra-Physics, CA) were used for the excitation of cyXR w/wo lutein. The transient absorption (TA) spectra were obtained by monitoring the intensity change of white light from a Xe-arc lamp (L9289-01, Hamamatsu Photonics, Japan) passed through the sample using an ICCD linear array detector (C8808-01, Hamamatsu Photonics, Japan). To increase the signal-to-noise ratio, 40–60 spectra were averaged, and the singular-value decomposition analysis was applied. To measure the time evolution of transient absorption change at specific wavelengths, the output of a Xe-arc lamp (L9289-01, Hamamatsu Photonics, Japan) was monochromated using monochromators (S-10, Soma Optics, Japan), and the change in the intensity after the photoexcitation was monitored using a photomultiplier tube (R10699, Hamamatsu Photonics, Japan). To increase signal-to-noise ratio, 200–400 signals were averaged. Global fitting of the signals using a multi-exponential function was performed to determine the lifetimes and absorption spectra of each photointermediate.

To measure the fold change in transient absorption change between cyXR with and without lutein, nanosecond laser pulses from an optical parametric oscillator (basiScan, Spectra-Physics, CA) pumped by the third harmonics of Nd-YAG laser (*λ* = 355 nm, INDI40, Spectra-Physics, CA) were used for the excitation of cyXR at different wavelengths (*λ*_exc_ = 430, 455, 470, 485, 520, 545 and 585 nm). The pulse energy was adjusted to 0.55 mJ cm^−2^, which falls within the range that maintains the linearity between the number of the absorbed photons and the transient absorption change. To calculate the quantum efficiency of excitation energy transfer (𝛟_EET_) (see table percentages on Fig. 5B), sample solutions of rhodopsins, one without a bound lutein and the other with a bound lutein, and their absorbances at the excitation wavelengths, *Abs0* and *Abs*, respectively, were considered. The number of photons absorbed by the retinal in the rhodopsin without a bound lutein, 𝑁^0^ is expressed as:

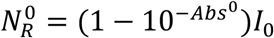

where 𝐼_0_ is the intensity of the incident light.

Assuming than the numbers of photons absorbed by the retinal and the lutein in the rhodopsin with a bound lutein, 𝑁_𝑅_ and 𝑁_𝐿_, respectively, are proportional to their absorption cross-sections, 𝑁_𝑅_ and 𝑁_𝐿_ can be expressed as:

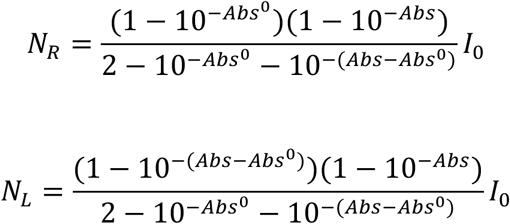

The ratio between the transient absorption signals of the rhodopsin samples without and with lutein, Δ𝐴𝑏𝑠^0^ and Δ𝐴𝑏𝑠, respectively, can be expressed as:

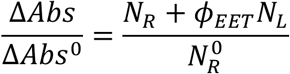

The 𝜙_𝐸𝐸𝑇_ can be expressed as follows using 𝑁^0^, 𝑁_𝑅_, 𝑁_𝐿_, Δ𝐴𝑏𝑠^0^, andΔ_𝐴𝑏𝑠_, which can be experimentally determined:

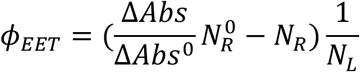

### Protein expression and purification for structural analysis

pBAD-cyXR was transfected in *E. coli* C41 (Rosetta). The transformant was grown in LB supplemented with 50 µg ml^−1^ ampicillin and 10 µg ml^−1^ at 220 r.p.m. at 37 °C. When the OD_600_ reached 0.6, expression was induced using a 0.1% final concentration of ʟ- arabinose (A3256, Sigma-Aldrich). The induced culture was grown at 120 r.p.m. overnight (more than 16 h) at 25 °C. Then, the pooled plate content was incubated with 20 µM all-trans retinal for more than 4 h in the dark. The collected cells were disrupted by sonication in buffer containing 20 mM Tris-HCl pH 8.0, 150 mM NaCl and 10% glycerol. The crude membrane fraction was collected by ultracentrifugation at 180,000g for 1 h. The membrane fraction was solubilized in buffer containing 20 mM Tris-HCl pH 8.0, 150 mM NaCl, 1% DDM and 10% glycerol for 2 h at 4 °C. The supernatant was separated from the insoluble material by ultracentrifugation at 180,000g for 20 min and incubated with Ni-NTA resin (Qiagen) for 30 min. The resin was washed with 10 column volumes of wash buffer containing 20 mM Tris-HCl pH 8.0, 500 mM NaCl, 0.03% DDM, 10% glycerol and 20 mM imidazole. The protein was eluted in buffer containing 20 mM Tris-HCl pH 8.0, 150 mM NaCl, 0.03% DDM, 10% glycerol and 300 mM imidazole. Eluted cyXR was concentrated and purified through size-exclusion chromatography on a Superose 6 Increase 10/300 GL column (GE Healthcare) with buffer containing 20 mM Tris-NaCl pH 8.0, 150 mM NaCl and 0.03% DDM. The protein was incubated with a buffer containing 20 mM Tris-NaCl pH8.0, 150 mM NaCl, 0.03% DDM and 1.75 mM lutein.

### XR structure predictions

For assessment of the presence of fenestrations in XRs for which no experimental structures were available, the structures were predicted with AlphaFold 3 with retinal as a covalent modification and 20 random seeds. PyMOL v. 3.1.3.1^38^ application programming interface and graphical user interface with the default solvent settings were used to screen the structures. To visualize the structural prediction of cyXR, the low-quality N- and C-terminal extensions were trimmed to exclude terminal residues with the per-residue average pLDDT (predicted local distance difference test) score of backbone atoms below 80.

### Cryo-EM single-particle analysis of the cyXR-lutein complex

For the cryo-EM grid preparation of the lutein-bound cyXR, the protein was reconstituted in nanodiscs. cyXR, MSP2N2, and SoyPC were mixed at a molar ratio of 1:4:200, respectively, and rotated at 4 2 for 1 h. Detergents were removed by adding Bio-Beads SM2 (Bio-Rad) to 40 mg ml^-1^, followed by gentle agitation. The Bio-Beads (equal amount) were incubated at 4 2 overnight. The Bio-Beads were then removed, and the solution was ultracentrifuged before size-exclusion chromatography.

The ultracentrifuged sample was concentrated and purified through size-exclusion chromatography on a Superose 6 Increase 10/300 GL column (GE Healthcare) with buffer containing 20 mM Tris-NaCl pH8.0 and 150 mM NaCl. The peak fractions of the protein were collected and concentrated to an absorbance (A280) of 15, using a centrifugal filter unit (50 kDa molecular weight cut-off; Merck Millipore). Of protein, 3 µl was loaded onto glow-discharged holey carbon grids (Quantifoil Au 300 mesh R1.2/1.3), after which, these were plunge-frozen in liquid ethane, using a Vitrobot Mark IV (Thermo Fisher Scientific). Cryo-EM imaging data were collected on a Titan Krios at 300 kV, equipped with a Gatan K3 Summit detector and controlled using the EPU software (Thermo Fisher’s single-particle data collection software). Images were obtained at a dose rate of about 15.5940 e^−^ Å^−2^ s^−1^, with a defocus ranging from −0.6 to −1.6 μm. Total exposure time was 2.3 s, with 48 frames recorded per micrograph. A total of 12,126 videos were collected. All acquired movies were dose-fractionated and subjected to beam-induced motion correction implemented in cryoSPARC^39^. The contrast transfer function (CTF) parameters were estimated using cryoSPARC^40^. Particles were initially picked from a small fraction with Gaussian blob picking and subjected to 2D classification. Class averages showing reasonable features of the cyXR pentamer in various orientations were selected as templates for template-based particle picking. Particles from these class averages generated an ab initio model in cryoSPARC. For each full dataset, extracted particles were downsampled to 3.32 Å, followed by two rounds of 2D classification to remove ‘junk’ particles. 3D classification into two classes with the ab initio model as a reference was performed using C5 symmetry. After multiple rounds of 3D classification, particles were re-extracted with the pixel size of 1.10 Å and a box size of 288 pixels. Multiple rounds of local CTF refinement and non-uniform refinement^41^ were performed using cryoSPARC. Finally, the 327,293 particles in the best class were reconstructed using non-uniform refinement, resulting in a 2.64 Å resolution reconstruction, with the gold-standard Fourier shell correlation (FSC = 0.143) criteria^42^. The quality of the map was sufficient to build a model manually in Coot^43,44^.

Model building was performed based on the initial model generated by ModelAngelo^45^ from the amino acid sequence of cyXR. We manually readjusted the model into the density map using Coot and refined it using phenix.real_space_refine (v.1.19)^46,47^. The asymmetric lutein molecule was tested in two orientations due to the lack of clear density to determine its precise orientation. The final placement was selected based on the orientation that best fit the density map.

## Results

### Phylogenetic analysis of XRs in cyanobacteria

Our search for rhodopsin genes in cyanobacterial genomes revealed that 11.7% and 14.7% of representative genomes possess them in GTDB and JGI GEM, respectively, with XRs present in 14.1%/21.5% of rhodopsin-carrying genomes-surpassed by bacterial halorhodopsins (anion pumps) and members of the xenorhodopsin family (inward proton pumps and sensory rhodopsins) (Supplementary Fig. S1). Despite these modest numbers, XR genes were detected in diverse and often unrelated cyanobacterial families. To our surprise, the majority of the cyanobacterial XRs came from families of filamentous cyanobacteria, not only MAGs but also isolates of *Nodosilinea antarctica* (non-axenic cultures), *Halomicronema* spp., *Phormidesmis priestleyi*, *Leptothoe* sp. (*Leptolyngbya* sp. Heron Island J), *Leptolyngbyaceae* sp. JSC-12 and *Rivularia radiosa* (*Plectonema* sp. *radiosum* LEGE 06105). Interestingly, all of the XRs from cyanobacterial genomes appeared to cluster together in a single clade (Fig. 1A). This clade was nested among XR sequences originating from *Deinococcota* with good support thus indicating that cyanobacteria acquired the first XR genes from this extremophile group.

A fine-grained phylogenetic analysis of cyanobacterial XRs together with related sequences from unbinned environmental contigs, showed that XRs coming from genomes assigned to the same cyanobacterial family often do not form a holophyletic grouping as is the case with the genes from the *Chamaesiphonaceae*, *Elainellaceae*, *Leptolyngbyaceae*, *Nostocaceae*, *Phormidesmiaceae* and *Pseudanabaenaceae* which further indicates that horizontal gene transfer between cyanobacterial groups played a major in spreading the XR genes among them (Fig. 1B). In this analysis, the cyanobacterial clade is further subdivided into four well-supported subclades (A−D), each with its environmental and taxonomic trends. The largest subclade, A, is composed of XR genes from unrelated cyanobacterial families with dissimilar morphotypes ranging from the unicellular thylakoid-less *G. violaceus* (the well-characterized GR) to the family of the epiphytic *Chamaesiphonaceae* and filamentous families (including cyXR characterized here). This subclade covers sequences originating from diverse terrestrial, freshwater, and saline habitats. Most conspicuously, all XR genes from cold environments: glaciers, meltwater, cryoconite and polar deserts, belong to this subclade. Further, considering that sequences from temperate locations, such as GR, constitute a minority of this subclade, this strongly suggests that subclade A underwent adaptation to low temperatures. The smaller subclades B, C and D were found in isolates and environmental datasets originating strictly from aquatic habitats from temperate waters or hot springs and lakes. The cyanobacteria possessing these genes, when known or predictable, are filamentous, except for *Halothece* spp. which possess subclade-B XR genes.

The genomic context of the XR genes in different cyanobacteria was found to be highly diverse with no indication that any of them belong to an operon. Thus, no *blh* genes for retinal synthesis from beta-carotene typically encountered in the same operon as proteorhodopsin and xanthorhodopsin genes in diverse bacteria^48,49^ or any other gene for carotenoid biosynthesis were found in the vicinity of the cyanobacterial XR genes (Supplementary Fig. S2). In fact, none of the XR-possessing cyanobacterial genome assemblies were found to have the *blh* gene in any other genomic location either. The most frequent gene neighbors of XR genes instead were widespread cyanobacterial gene families *hli* coding for high-light inducible proteins (*e.g*., upstream of the GR gene) and *nblA* coding for the proteolysis adaptor NblA, in the vicinity of XR genes from subclades A and D. While light-dependent expression of *hli*^50^ and XR genes, as exemplified by GR^51^, might explain their genomic proximity, no functional connection to *nblA* is apparent.

Given the recently reported gene regulatory activity of GR^51^, we also assessed the appearance of genes coding for ArsR-like helix-turn-helix transcription regulators similar to GvTcR from *G. violaceus* in XR-possessing cyanobacteria. Only genomes from the *Elainellaceae* genus JAAUTN01 were found to have related genes (≥50% identity to GvTcR) indicating that the interaction between GR and GvTcR might be specific to *G. violaceus*.

The finding of XR genes in filamentous cyanobacteria is noteworthy. cyXR is thus only the second characterized example of an ion-pumping rhodopsin from filamentous bacteria following the discovery of the bacterial halorhodopsin *Mr*HR (= MastR) in the cyanobacterium *Mastigocladopsis repens* PCC 10914^52–54^.

All of the cyanobacterial XRs were found to invariably possess glycine at the fenestration-enabling position G156 and their predicted structures demonstrated a fenestration similar to that of XR, GR and Kin4B8-XR hinting at the possibility that not only GR but many other cyanobacterial XR might be able to bind carotenoid antennas as well.

### Functional characterization of cyXR

To investigate the ion transport function of cyXR, the protein was expressed in *E. coli* cells. The transformed cyXR-expressing *E. coli* exhibited a purple color, indicating the formation of a functional rhodopsin (Fig. 2A). The proton transport activity of cyXR in the *E. coli* cells was assayed by monitoring the light-induced changes in pH of the external solvent, which was used to characterize the functions of other microbial rhodopsins, like outward and inward proton, inward Cl^-^, and outward Na^+^ pumps^55–57^. The cell suspension of *E. coli* expressing rhodopsin proton pumps shows acidification (pH decrease) or alkalinization (pH increase) of the external medium upon illumination by outward or inward proton transport, respectively, and the signals are eliminated in the presence of a protonophore, CCCP. We observed acidification of the external solvent, and it was largely eliminated by the addition of 10 μM CCCP, suggesting that cyXR functions as a light-driven outward proton pump (Fig. 2A). The large pH decrease observed for cyXR following the addition of CCCP is consistent with the fact that it poses a DTE motif (Fig .2B and upplementary Fig. S3), which is the most common among the outward proton pumping rhodopsins and is known for its effectiveness in achieving higher transport activity^1,56,58,59^.

Our experiments comparing the proton-pumping activities of GR and cyXR revealed striking differences in temperature dependence. When expressed in *E. coli* cells, these two phylogenetically related rhodopsins displayed different responses to temperature reduction (Fig. 2C and D). GR exhibited the expected enhancement of apparent proton-pumping activity as temperature decreased from 20°C to 5°C, consistent with previous observations using heterologous *E. coli* expression systems. This temperature-dependent increase in apparent activity (approximately 3-fold stronger signal at 5°C compared to 20°C) is primarily attributed to reduced activity of endogenous *E. coli* proton transporters at lower temperatures, which decreases the rate of passive proton backflow across the membrane. Remarkably, and contrary to predicted expectations, cyXR demonstrated temperature-independent initial proton-pumping rates across the tested temperature range (5°C, 12.5°C, and 20°C). As shown in Fig. 2D, the initial slopes of light-induced pH changes for cyXR remained nearly constant across all temperatures, while GR showed a clear negative correlation between temperature and initial pumping rate. This unexpected behavior suggests that cyXR’s intrinsic pumping activity might actually decrease at lower temperatures likely due to its high activation enthalpy barrier that renders the photocycle significantly slower under low-temperature conditions than that in GR. The reduction in the proton-pumping turnover rate of cyXR counterbalances the reduced activity of endogenous *E. coli* proton transporters that would otherwise lead to enhanced signals at lower temperatures.The contrasting temperature responses between GR and cyXR indicate potential adaptations in the molecular mechanism of cyXR. This finding is particularly intriguing given that some cold-adapted organisms maintain membrane fluidity through increased proportions of unsaturated fatty acids. The native lipid environment of cyXR likely differs significantly from that in the *E. coli* expression system, which may influence its temperature-dependent functionality.

To examine the *λ*_max_ and the expression level of cyXR, hydroxylamine (HA) bleaching assay was conducted after solubilization of *E. coli* membranes with detergent, DDM. HA hydrolyzes the protein-bound retinal chromophore into retinal-oxime, which will be released so that the *λ*_max_ of the original protein can be estimated without purification by calculating the difference absorption spectra between before and after the reaction with HA. Fig. 2E shows the result representing the photobleaching of cyXR in the presence of 500 mM HA. The positive peak at ∼542 nm corresponds to the absorption of the rhodopsin with the protonated retinal Schiff base (RSB) present prior to the bleaching. The small positive peak at ∼425 nm was attributed to the absorption of rhodopsin with the deprotonated RSB, while the negative peak at 361 nm was derived from the absorption of the reaction product, retinal-oxime. This result indicates that the expressed cyXR has the absorption spectrum mainly peaked in the visible region.

### Molecular properties and photocycle

To further study the molecular properties, we purified and spectroscopically investigated cyXR. The purified cyXR showed a *λ*_max_ at 545 nm in detergent (DDM) (Fig. 3A). The spectra of purified cyXR at different pH values are shown in Supplementary Fig. S4. The absorption peak in the visible wavelength region corresponding to the state with the deprotonated RSB, denoted by *λ*_max_ = 546 nm, decreased and slightly redshifted when the pH was raised, and simultaneously, another peak appeared in the ultra-violet (UV) region (*λ*_max_ = 366 nm) (Supplementary Fig. S4A, top). The difference absorbance values at these peak wavelengths were plotted against pH and fitted with the Henderson-Hasselbalch equation (Supplementary Fig. S4A, bottom), which determined that the RSB in cyXR exhibits two p*K*_a_ values of 10.8 ± 0.4 and 11.19 ± 0.02. These values were in line with the RSB p*K*_a_ of typical microbial rhodopsins (*e.g*., the RSB p*K*_a_ of bacteriorhodopsin is 13.3 ± 0.3)^60^. In contrast, a 13 nm red-shift in the *λ*_max_ was observed on the acidic side (Supplementary Fig. S4B, top). This is caused by the protonation of the counterion in the third transmembrane helix (TM3) (aspartic acid (D115), Fig. 2B), as known for many microbial rhodopsins^1,61–63^.

**Fig. 3.**
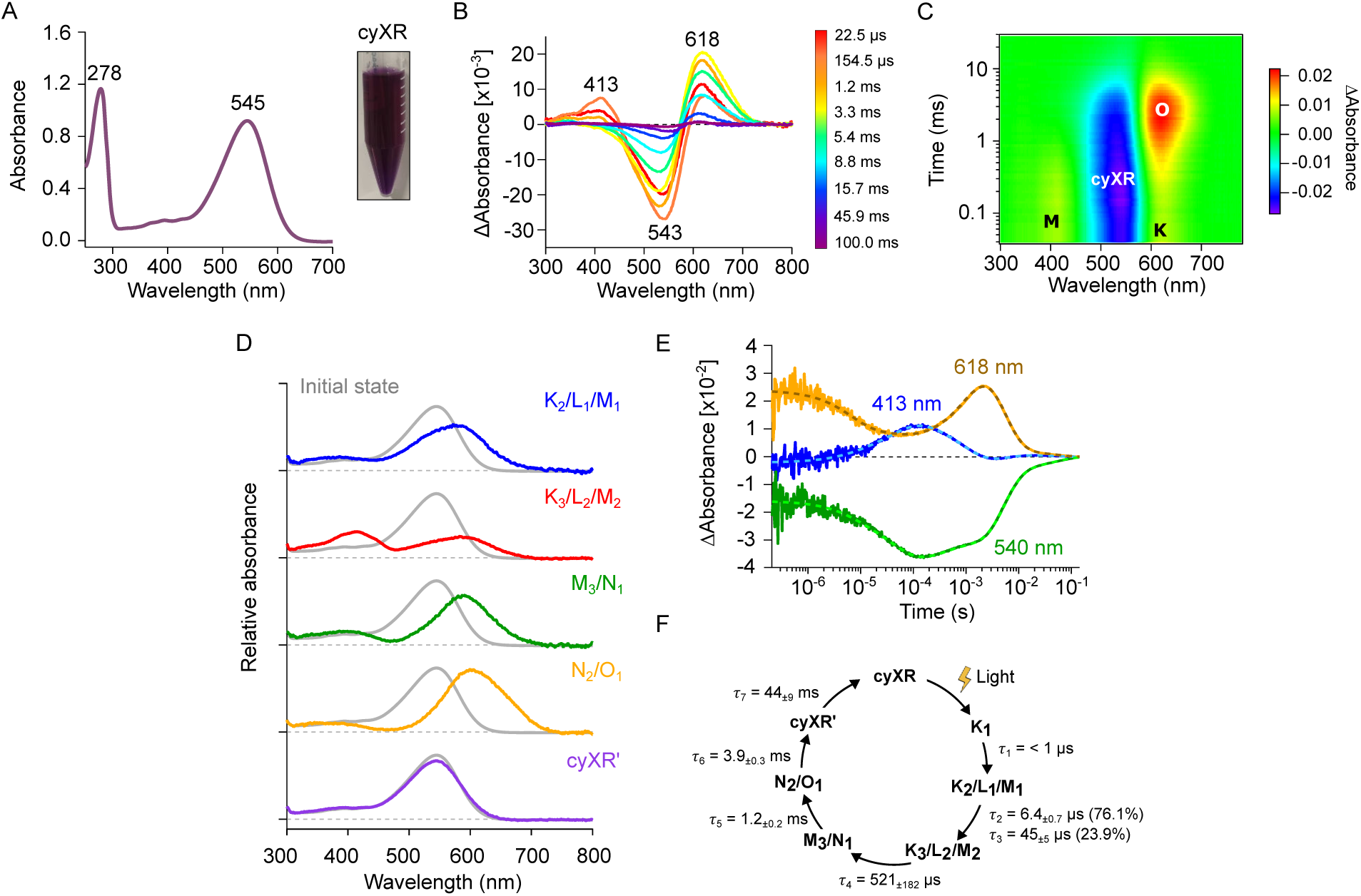
**Absorption spectra and photocycle of cyXR**. **A** UV-vis absorption spectrum for cyXR. The pictures of the purified proteins are shown next to the corresponding results; **B** Transient absorption spectra at different time points; **C** The two-dimensional plot of transient absorption change; **D** Absorption spectra of photointermediates. **E** Time course of the transient absorption change; **F** Photocycle model of cyXR.

**Fig. 4.**
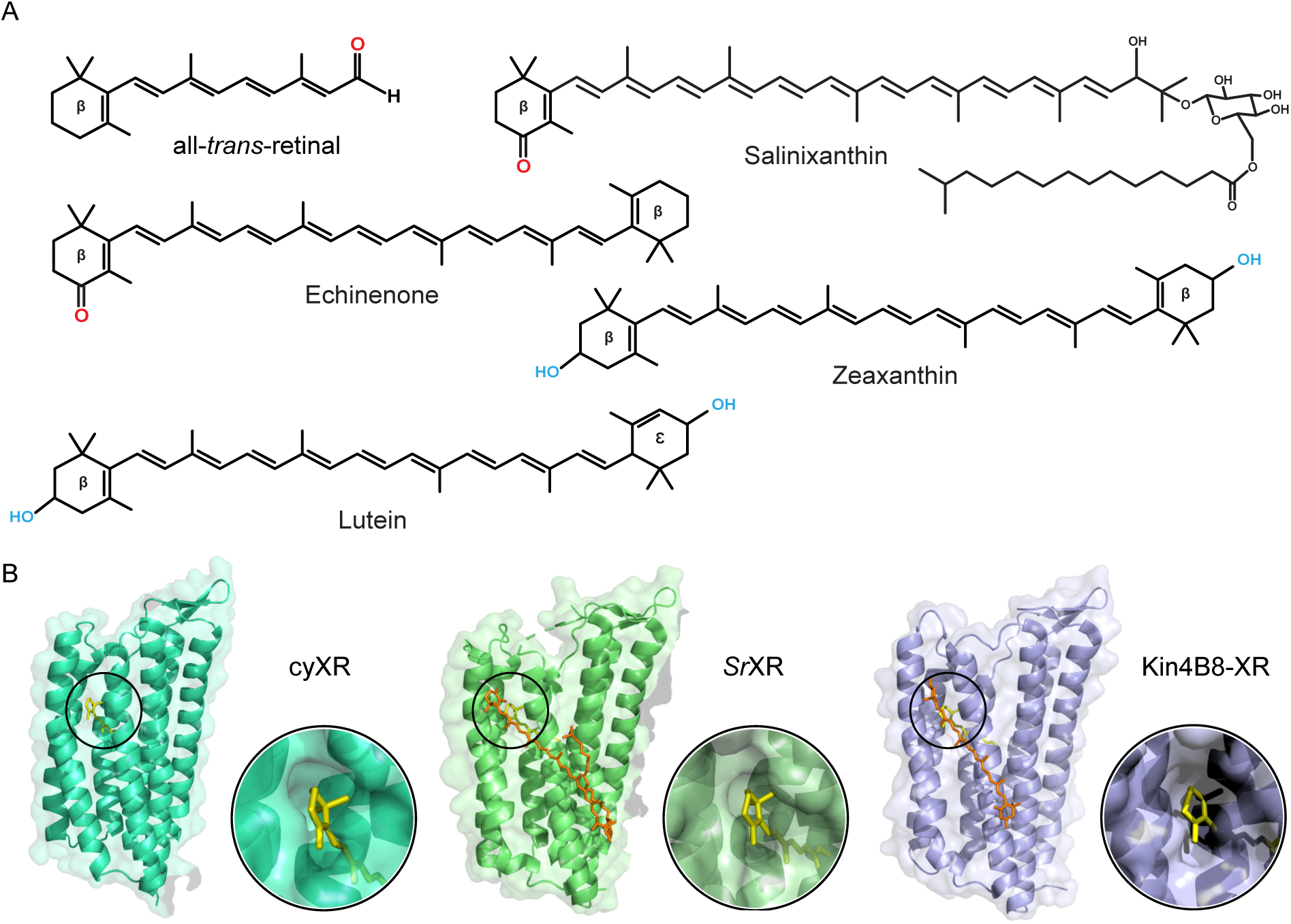
Carotenoids structures and XRs fenestrations. **A** The chemical structures of all-*trans*-retinal and various ketocarotenoids (salinixanthin and echinenone) and hydroxycarotenoids (zeaxanthin and lutein). The type of cyclic end groups (ꞵ and ε) of each carotenoid is also shown; **B** Structural comparison of fenestrated XRs: cyXR (AlphaFold3 model)^66^, *Sr*XR (PDB: 3DDL)^5^, and Kin4B8-XR (PDB: 8I2Z)^8^. Low- quality N- and C-terminal extensions are not shown (see Methods for details). The structures are aligned based on the retinal β-ionone ring position, and the fenestration zone is highlighted. For clarity, salinixanthin and zeaxanthin are not shown in the enlarged fenestration zone of *Sr*XR and Kin4B8-XR structures, respectively.

**Fig. 5.**
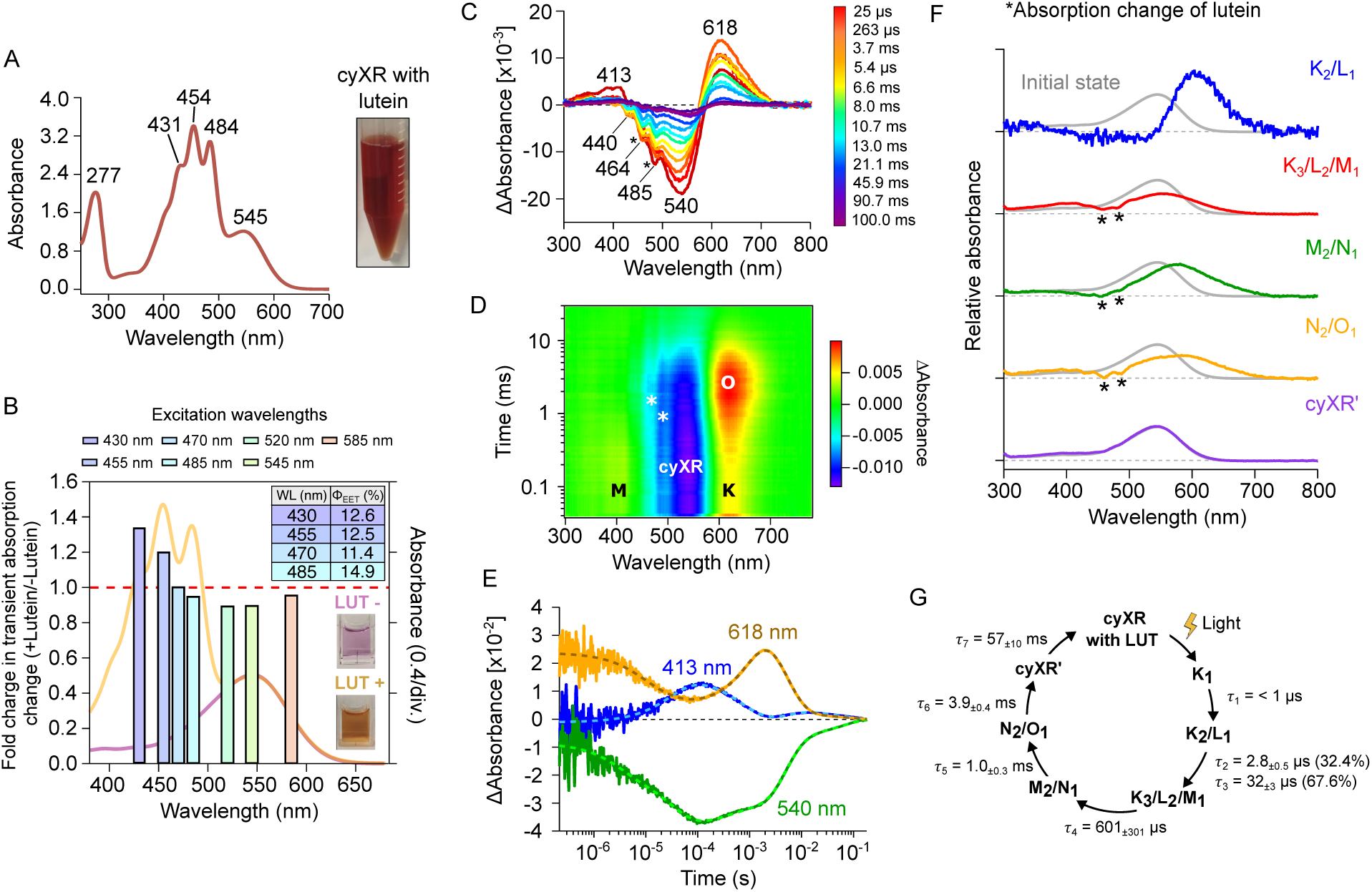
**Absorption spectra and photocycle of cyXR with lutein**. **A** UV–vis absorption spectrum of purified cyXR with lutein. The picture of the purified protein is also shown; **B** Ratios of transient absorption change in cyXR with and without lutein at different excitation wavelengths. Bars are colored according to the color of excitation light. Raw data of transient absorption spectra at each excitation wavelength are displayed in Fig. S7. The absorption spectra of cyXR without (purple line) and with (orange line) lutein were overlaid. The red dashed line indicates no difference between with and without lutein. Table with the 𝛟_EET_ percentages and the pictures of the purified proteins are shown next to the corresponding results; **C** Transient absorption spectra at different time points; **D** The two-dimensional plot of transient absorption change; **E** Time course of the transient absorption change; **F** Absorption spectra of photointermediates. Asterisks represent the absorption change of lutein; **G** Photocycle model of cyXR with lutein.

The retinal configuration was also investigated using high-performance liquid chromatography (HPLC) on retinal oximes produced by the hydrolysis reaction of the RSB in cyXR with HA (Supplementary Fig. S5). Most retinal chromophores derived from dark-adapted (DA) proteins showed an all-*trans* configuration with trace amounts of 13-*cis* and 11-*cis* ones (Supplementary Fig. S5A). During illumination at *λ* = 550 ± 10 nm, the 13-*cis*- retinal form accumulated substantially, while the amount of 11-*cis*-retinal form slightly increased (yellow bars in Supplementary Fig. S5B and Table S1). The all-*trans* form represents the functional state of cyXR, similar to typical microbial rhodopsins^2,64^.

Next, the photocycle reaction of cyXR was studied by laser-flash photolysis method, in which cyXR protein was solubilized in DDM. Fig. 3B shows the transient change in absorption of cyXR upon excitation at *λ*_max_ = 550 nm, representing the accumulation of blue-shifted (M) and red-shifted (K and O) photointermediates (Fig. 3C) at 413 nm and 618 nm, respectively, in addition to L, N, and cyXR’ intermediates with *λ*_max_ close to the initial state (Fig. 3D). Simultaneously, the maximum bleaching signal at 543 mn was observed, indicating the M intermediate accumulation at *t* ∼155 us. To monitor the transient absorption change representing the short-lived photointermediates prior to M, we conducted measurement at specific probe wavelengths using a photomultiplier tube at higher time resolution (Fig. 3E). The photocycle and the absorption spectra of the initial state and five photointermediates (K_2_/L_1_/M_1_, K_3_/L_2_/M_2_, M_3_/N_1_, N_2_/O_1_, and cyXR’) were determined by global fitting analysis of the transient absorption changes (Fig. 3F).

### Hydroxycarotenoid-binding to cyXR

The ability to bind a carotenoid antenna in rhodopsin proteins requires the presence of a fenestration, an opening in the protein moiety that exposes the retinal-binding pocket to the external environment. This structural feature facilitates energy transfer between the two chromophores and is determined by a conserved glycine residue at a specific position in the amino acid sequence (see Supplementary Fig. S3, black circle)^6^. Collected XRs (as well as PRs) exhibit variability at this position, with some representatives possessing a bulky tryptophan or phenylalanine that precludes carotenoid antenna binding, while others maintain the glycine residue, allowing for such binding^8,65^. However, there is limited evidence of other XRs with this specific glycine residue using carotenoid antennas in nature.

We hypothesized that cyXR could potentially utilize a ketolated carotenoid as a light-harvesting antenna, as observed in previous energy transfer studies involving light-harvesting ketocarotenoids and light-driven proton pump XRs in bacterial systems such as *Sr*XR with salinixanthin^5,6^ and GR with echinenone^9^ (Fig. 4A). As shown in Fig. 4B, cyXR is predicted to contain a fenestration exposing the retinal ring, similar to *Sr*XR^5^ and, as a member of the XR family, it contains a glycine residue at the *Sr*XR position 156 in TM5 (G115 in cyXR, Fig. 2B). To investigate the impact of carotenoid antenna binding on the properties of the retinal binding pocket, we attempted to purify cyXR with a commercially available ketolated carotenoid pigment, canthaxanthin, which is widely distributed in nature (Supplementary Fig. S6A). Our findings revealed that canthaxanthin, despite being a ketolated carotenoid, did not bind to cyXR (Supplementary Fig. S6B).

Motivated by this result, we explored the binding potential of cyXR with a hydroxylated carotenoid, as observed in other bacterial XRs, such as Kin4B8-XR^8^ with lutein and zeaxanthin (see Fig. 4A). The chosen candidate was the commercially available hydroxylated carotenoid lutein (Fig. 4A and Supplementary Fig. S6C). Incubation of purified cyXR with lutein resulted in binding (Fig. 5A). Energy transfer from lutein to the retinal molecule of cyXR was investigated by laser-flash photolysis (Fig. 5B).

It revealed an enhancement in the transient absorption (TA) signal for cyXR bound to lutein when excited at the 450–490 nm region, supporting the energy transfer from the carotenoid to the retinal, enhancing the retinal isomerization. cyXR photocycle was enhanced by 1.3 folds when bound to lutein, which is similar compared to the value of Kin4B8-XR and a little more than half compared to the value of HeimdallR1 (1.6 and 3.2 folds enhancement, respectively)^8,10^ bound to lutein. The excitation energy transfer quantum yield (𝜙_𝐸𝐸𝑇_) between the retinal and lutein chromophores was estimated to be more than 10% and nearly constant (see table of percentages values on Fig. 5B) over all excitation wavelengths.

To investigate how the retinal binding pocket is affected by lutein binding, we compared the absorption of purified cyXR with and without lutein (Figs. 3A and 5A). Binding lutein did not display a shift in the observed *λ*_max_ when compared with cyXR alone, which was also observed in bacterial rhodopsins investigated previously^6,8^. The absorption peak of the retinal was decreased and slightly red-shifted when the pH was raised, corresponding to the structural change affecting the deprotonated RSB in cyXR, reproducing the behavior observed in cyXR alone (Supplementary Fig. S4A, top). In the case of the acidic side, a red-shift in the *λ*_max_ representing the protonation state of the counterion was observed as cyXR alone (Supplementary Fig. S4B, bottom). Hence no difference in the *λ*_max_ was observed between cyXR with and without lutein, indicating the impact of the lutein binding on the property of the retinal binding site is minor. The influence of the lutein binding on the retinal isomer composition in cyXR was also investigated using HPLC of retinal oximes produced by hydrolyzing the RSB with HA (Supplementary Fig. S5B). The binding of lutein increased the fraction of the all-*trans* form in the dark from 92.6% to 95.5%, which represents the functional form to initiate proton-pumping photocycle (Supplementary Fig. S5B, bottom and Table S1). The transient absorption changes observed for cyXR with lutein are roughly similar to those observed for cyXR without lutein (Figs. 5C−F). Multi-exponential analysis identified 5 photointermediates having different absorption spectra (K_2_/L_1_, K_3_/L_2_/M_1_, M_2_/N_1_, N_2_/O_1_, and cyXR’) (Figs. 5F and G) . Two additional sharp peaks around 465 nm and 485 nm (see asterisks in Figs. 5C, D and F), which were absent for cyXR alone, were observed when lutein was bound to the protein, indicating that a conformational change of the protein occurs during the photocycle that alters the structure and the absorption of lutein. The observed biphasic spectral feature upon carotenoid antenna binding to the protein, characterized by two distinct sharp peaks, contrasts with the monophasic spectral profile, exhibiting a single sharp peak, previously reported for Kin4B8-XR^8,10^.

### Overall structure of cyXR with lutein

To investigate the xanthophyll-binding properties of cyXR, we determined a cryo-electron microscopy structure of the lutein-bound cyXR at a nominal resolution of 2.64 Å (Supplementary Figs. S8A−C and Table S2). cyXR assembles into a pentameric configuration (Fig. 6A), similar to Kin4B8-XR (Protein Data Bank ID: 8I2Z)^8^. The monomeric units of cyXR exhibit highly conserved features with other XRs (Kin4B8-XR and *Sr*XR)^67^, including similar carotenoid-binding orientations (Supplementary Fig. S8D).

**Fig. 6.**
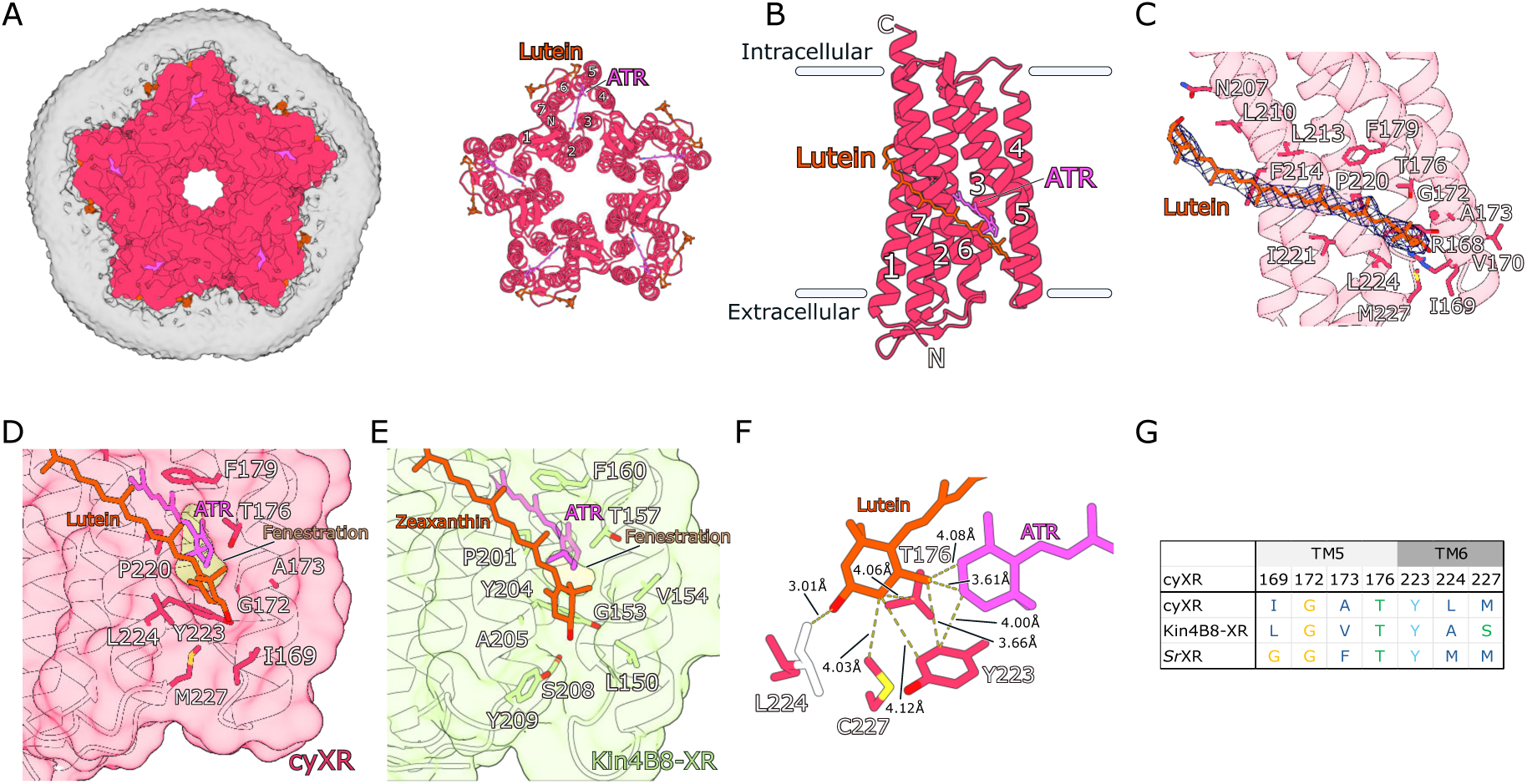
**Structure of cyXR bound to lutein**. **A** Electron microscopy map and pentameric structure of the lutein-bound cyXR, viewed from the extracellular side. Lutein and all-*trans* retinal (ATR) are shown as stick models; **B** Overall structure of the monomeric unit, with lutein and the retinal chromophores; **C** Lutein-binding site with the cryo-EM density. The extended carotenoid is tightly bound to the transmembrane surface of cyXR, traversing nearly the entire bilayer, with an inclination of about 60° to the membrane normal; **D** and **E** Fenestrations in cyXR (D) and Kin4B8-XR (E); **F** Positional relationship between the hydroxyl ring of lutein, retinal and surrounding residues; **G** Conservation of the residues surrounding the fenestrations in XRs. Glycine is coloured orange. Polar and hydrophobic residues are coloured green and blue. Tyrosine is coloured cyan.

In cyXR, lutein is positioned transversely along the outer surface of TM6 (Fig. 6C), with its hydroxyl ring fitting into a fenestration of the protein. Notably, this fenestration contains a G172 residue, which corresponds to G153 in Kin4B8-XR, highlighting a structural conservation between the two fenestrations. This binding mode closely resembles that of zeaxanthin in Kin4B8-XR (Figs. 6D and E), suggesting a conserved mechanism of energy transfer. However, unlike Kin4B8-XR, the fenestration at the base of cyXR is less hydrophilic, although the backbone carbonyl group of I169 may still form a hydrogen bond with lutein. Furthermore, the residue corresponding to A205 in Kin4B8- XR is replaced by L224 in cyXR, whose bulkier side chain imposes a steric constraint. This constraint slightly shifts the orientation of lutein’s ring, resulting in a distinct arrangement compared to zeaxanthin in Kin4B8-XR (Fig. 6 F and G; Supplementary Fig. S8E). Such structural features of the fenestration may contribute to maintaining a stable energy transfer mechanism by reducing the distance between the lutein ring and the *β*- ionone ring of the retinal, while also preventing the intrusion of water molecules into the fenestration.

## Discussion

Our study presents the discovery and characterization of a novel xanthorhodopsin (cyXR) in an Antarctic cyanobacterium, expanding our understanding of microbial rhodopsins in these important photosynthetic organisms. This finding bridges a significant gap in our knowledge of XRs in cyanobacteria and provides new insights into their adaptations to diverse light environments.

The phylogenetic analysis reveals that cyXR belongs to a distinct clade of XRs found predominantly in cold environments, including polar regions. This suggests a potential adaptive role of these proteins in low-temperature habitats. Notably, cyXR represents the first example of XRs in filamentous microorganisms, expanding our understanding of rhodopsin distribution and evolution in more complex microbial life forms. The presence of cyXR in Antarctic cyanobacteria suggests an important role in their adaptation to extreme environments. Unlike the previously described dual systems of XRs and anoxygenic photosynthesis, our findings indicate that cyXR complements oxygenic photosynthesis in Antarctic cyanobacteria. This dual strategy of utilizing both chlorophyll-based photosynthesis and rhodopsin-based phototrophy may provide these organisms with a competitive advantage in environments with variable light conditions.

Our functional characterization confirms that cyXR acts as a light-driven outward proton pump, as other XRs reported in literature. The absorption maximum at 545 nm in DDM detergent indicates that cyXR is well-suited for capturing light in the green-yellow part of the spectrum, potentially complementing the chlorophyll-based photosynthetic apparatus in these cyanobacteria. The detailed photocycle analysis of cyXR, both with and without lutein, provides insights into its functional mechanism. The observed enhancement of the photocycle by lutein binding, although modest, suggests a potential role in optimizing light capture under the variable and often low-light conditions of Antarctic ecosystems. The unique spectral changes observed during the photocycle of lutein-bound cyXR, particularly the two sharp peaks at 465 nm and 485 nm, indicate a novel protein-carotenoid interaction not previously observed in other bacterial rhodopsins.

Moreover, the most intriguing aspect of our findings is the unexpected binding of lutein, a hydroxylated carotenoid, to cyXR. This is in contrast to the previously reported example of carotenoid binding to a XR member in cyanobacteria where a ketolated carotenoids (salinixanthin, echinenone and canthaxanthin) bound GR. The energy transfer from lutein to the retinal chromophore, albeit with a lower efficiency compared to some other XRs, suggests a novel adaptation in light-harvesting strategies. This finding expands our understanding of the diversity of carotenoid-rhodopsin interactions and may indicate an evolutionary adaptation to the specific light conditions in Antarctic environments.

The discovery and characterization of cyXR not only expands our understanding of microbial rhodopsins in cyanobacteria, but also provides new insights into microbial adaptations to extreme environments. The unique ability of cyXR to bind a hydroxylated carotenoid, as opposed to the previously documented ketocarotenoid-binding GR, suggests a diversification in light-harvesting strategies among cyanobacteria. This adaptation likely plays a crucial role in helping cyanobacteria thrive in diverse and fluctuating light environments, contributing to their ecological success.

The cryo-electron microscopy structure of lutein-bound cyXR provides detailed insights into the xanthophyll-binding mechanism of this rhodopsin variant. The lutein-binding fenestration in cyXR induces a rotation of the carotenoid’s *β*-ionone ring: residue L224 protrudes into the pocket and pushes the ring upward (Fig. 6D and supplementary Fig. S8E). This rotation shortens the effective conjugation length and accounts for the uniform ∼3 nm hypsochromic shift of the three lutein absorption peaks (431, 454, 484 nm) relative to those reported for Kin4B8-XR^8^. Because snow and ice preferentially remove longer wavelengths, the ambient light spectrum is enriched in shorter wavelengths, making this blue-shift a plausible spectral adaptation. The same steric effect also narrows the distance between the lutein and retinal ionone rings, a geometry expected to enhance energy transfer. Thus, a single pocket residue can simultaneously tune the antenna spectrum and improve chromophore coupling, underscoring the adaptive flexibility of xanthorhodopsins.

Moreover, the connection between rhodopsins and photosynthesis emphasizes the potential relationship between these mechanisms. While Kopejtka *et al*.^7^ described ‘dual phototrophy’^14^ in a bacterium performing anoxygenic photosynthesis that also uses XRs for proton-pumping, our findings suggest another type of ‘dual phototrophy’ where cyXR is used by cyanobacteria that perform oxygenic photosynthesis (see Fig. 7 for various extant types of phototrophy currently found on earth). This connection underscores the importance of XRs in optimizing energy capture and enhancing the overall efficiency of photosynthetic (anoxygenic and oxygenic) processes. In conclusion, this hydroxylated carotenoid-binding XR represents an advancement in our understanding of cyanobacterial adaptation and energy acquisition. Further research into the diversity and function of XRs in cyanobacteria might shed light on the evolutionary strategies that enable these organisms to dominate a wide range of aquatic environments.

**Fig. 7.**
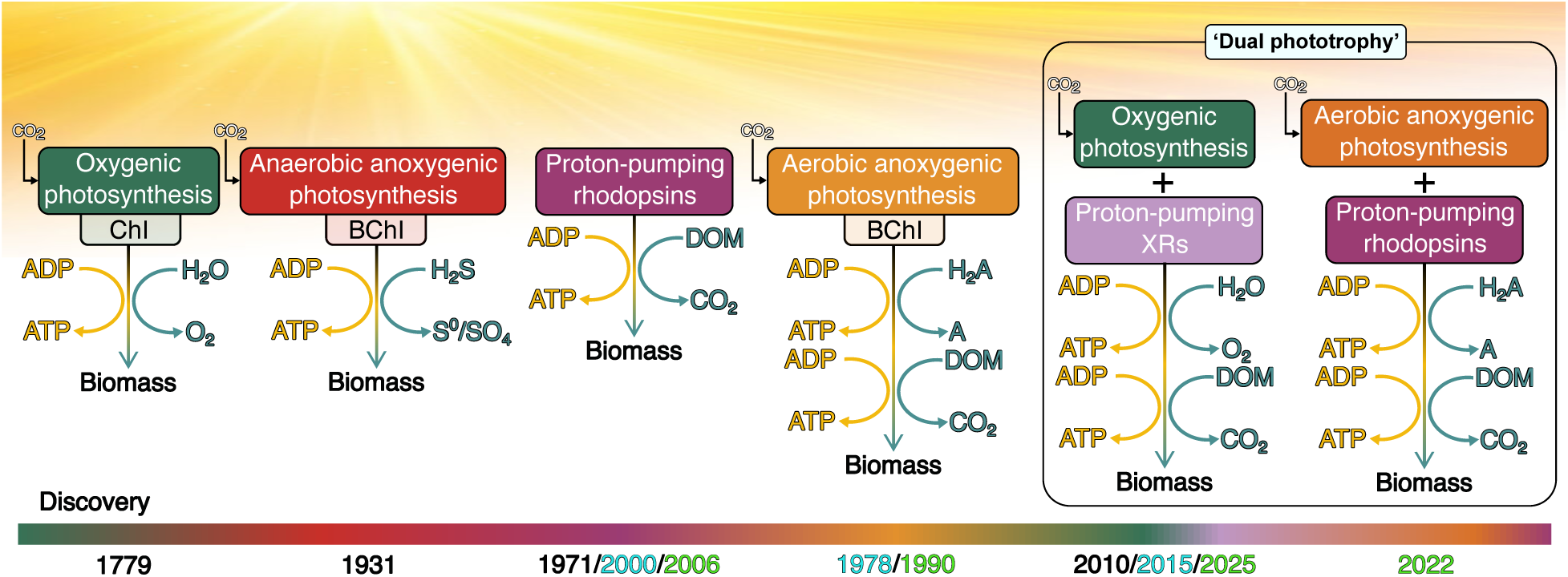
Diverse extant phototrophy mechanisms used by photo(hetero)trophic organisms. The timeline at the bottom highlights key discoveries of these processes and underscores the progressive understanding of solar energy utilization in different ecosystems, from oxygenic photosynthesis (1779) to ‘dual phototrophy’ (2022 & 2025). Years in black indicate the original description of the mechanism, turquoise denotes the first discovery in marine ecosystems, and green represents the first discovery in freshwater ecosystems. The figure was inspired by Karl (2002)^68^ and Sommaruga (2025)^14^.

## Data availability

Supplementary data and metadata for the bioinformatic pipeline are available from the Zenodo repository doi:10.5281/zenodo.15602260. The code used for the analyses is available from https://github.com/BejaLab/cyXR.

## Acknowledgments

M.d.C.M. is grateful to the Azrieli Foundation for the award of the Azrieli Fellowship (cohort 2022-2023) and its support during a 3-months research stay at the Institute for Solid State Physics (ISSP, University of Tokyo, Japan). M.d.C.M. also thanks the ISSP for its collaboration and support during this period. This work was supported by the Israel Science Foundation (Research Center grant 3131/20 to O.B.), the Nancy and Stephen Grand Technion Energy Program (GTEP), JSPS KAKENHI Grants-in-Aid (grants JP23H04404 to K.I., JP23K05007 to M.K., JP24K23232 to T.T., and JP24KJ0909 to S.M.), JST CREST (grants JPMJCR22N2 to K.I. and JPMJCR20E2 to O.N.), and MEXT

Promotion of Development of a Joint Usage/ Research System Project: Coalition of Universities for Research Excellence Program (CURE) (grant JPMXP1323015482 to K.I.), the Platform Project for Supporting Drug Discovery and Life Science Research (Basis for Supporting Innovative Drug Discovery and Life Science Research (BINDS)) from AMED, grant numbers JP24ama121002 (support number 3272, O.N.) and JP24ama121012 (O.N.), Research Foundation for Opto-Science and Technology (W.S.), and Brain Science Foundation (W.S.). O.B. holds the Louis and Lyra Richmond Chair in Life Sciences.

## Author contributions

M.d.C.M. and O.B. conceived the project. M.d.C.M. performed molecular biology, protein expression in *E. coli*, measurement of λ_max_, protein purification, purification of lutein-binding complex, HPLC analysis of retinal isomers, pH titration, experimental data analysis, and together with M.K. performed ion-pump assay measurements. M.d.C.M. and K.I. performed laser-flash photolysis. A.R. performed bioinformatics. S.M., T.T., W.S., and O.N. performed structural analysis. M.d.C.M., A.R., K.I., and O.B. wrote the paper with input from all authors.

