## Supplementary material for "Light-harvesting by antenna-containing xanthorhodopsin from an Antarctic cyanobacterium": Suplementary Materials

[1]Faculty of Biology, Technion-Israel Institute of Technology, Haifa 3200003, Israel. [2]Department of Biological Sciences, Graduate School of Science, The University of Tokyo, Tokyo, 113-0033 , Japan. [3]The Institute for Solid State Physics, The University of Tokyo, Kashiwa, 277-8581, Chiba, Japan. [4]The Nancy and Stephen Grand Technion Energy Program (GTEP), Technion-Israel Institute of Technology, Haifa 3200003, Israel. [5]Current address: Physical and Analytical Chemistry Department, University of Jaén, Jaén, Spain. [6]Current address: Department of Signal Exploration, The Sakaguchi Laboratory, Keio University School of Medicine, Tokyo, 160-8582, Japan. [7]These authors contributed equally: María del Carmen Marín, Shunya Murakoshi. *

**Amino acid sequence of cyXR**

>3300012044.a:Ga0136636_10013288_XR_OTU-4691
MSISLITDFALLAQSNLPETVSKVDFLSSGQYNLVYNAFSFAIATMLAAALFFFNVRG
QVGQKYQLALIVSALVVSIAGYHYYRIFGSWEAAYSLQNGNYVLTGAPFNDAYRYVD
WLLTVPLLLVETVAVLALPGMEARPLLIKLVVAAILMIATGYPGEISTDLNTRIVWGAVS
TVPFIYILYILWIELSKSLLRQPEGVQTLVKNMRLLLLFSWGVYPIAYLLPMLGISGATA
DVGVQIGYTIADVLAKPIFGLLVFAIALEKTRVDAGEQPTKESPMTMISS

**Supplementary Table S1. Retinal configuration in cyXR without and with lutein.** The composition of the retinal isomers was determined through HPLC analysis for retinal oxime produced by the hydrolysis of the retinylidene Schiff base with hydroxylamine. AT, 13C, and 11C indicate all-*trans*, 13-*cis*, and 11-*cis* configurations, respectively.

| Protein | Light conditions | AT (%) | 13C (%) | 11C (%) |
|---|---|---|---|---|
| cyXR | DA | 92.6 ± 0.9 | 6.0 ± 0.8 | 1.4 ± 0.2 |
| | Light | 74.9 ± 0.9 | 23.0 ± 0.7 | 2.1 ± 0.2 |
| | LA | 90 ± 1 | 8 ± 1 | 1.6 ± 0.4 |
| cyXR with lutein | DA | 95.5 ± 0.2 | 4.1 ± 0.2 | 0.40 ± 0.02 |
| | Light | 80.5 ± 0.4 | 17.9 ± 0.2 | 1.5 ± 0.2 |
| | LA | 93.91 ± 0.05 | 5.49 ± 0.05 | 0.59 ± 0.01 |

**Supplementary Table S2. Data collection, refinement and validation statistics for the cryo-EM structure of the cyXR-lutein complex.**

| | cyXR with lutein |
|---|---|
| **Data collection and procesing** | |
| Microscope | Titan Krios G3i |
| Magnification | 105,000 |
| Voltage (kV) | 300 |
| Electron expodure (e-/A2) | 49.3 |
| Defocus range (um) | -0.6 to -1.6 |
| Pixcel size (A/px) | 0.83 |
| Symmetry imposed | C5 |
| Initial particle images (no.) | 7,937,467 |
| Final particle images (no.) | 327,293 |
| Map resolution (Å) | 2.64 |
| FSC threshold | 0.143 |
| **Refinement** | |
| Atoms | 1,970 |
| R.m.s. deviations | |
| Bond lengths (Å) | 0.003 |
| Bond angles (°) | 0.57 |
| Validation | |
| Clashcore | 4.62 |
| Rotamers (%) | 3.12 |
| Ramachandran plot | |
| Favored (%) | 99.01 |
| Allowed (%) | 0.99 |
| Outliers (%) | 0 |

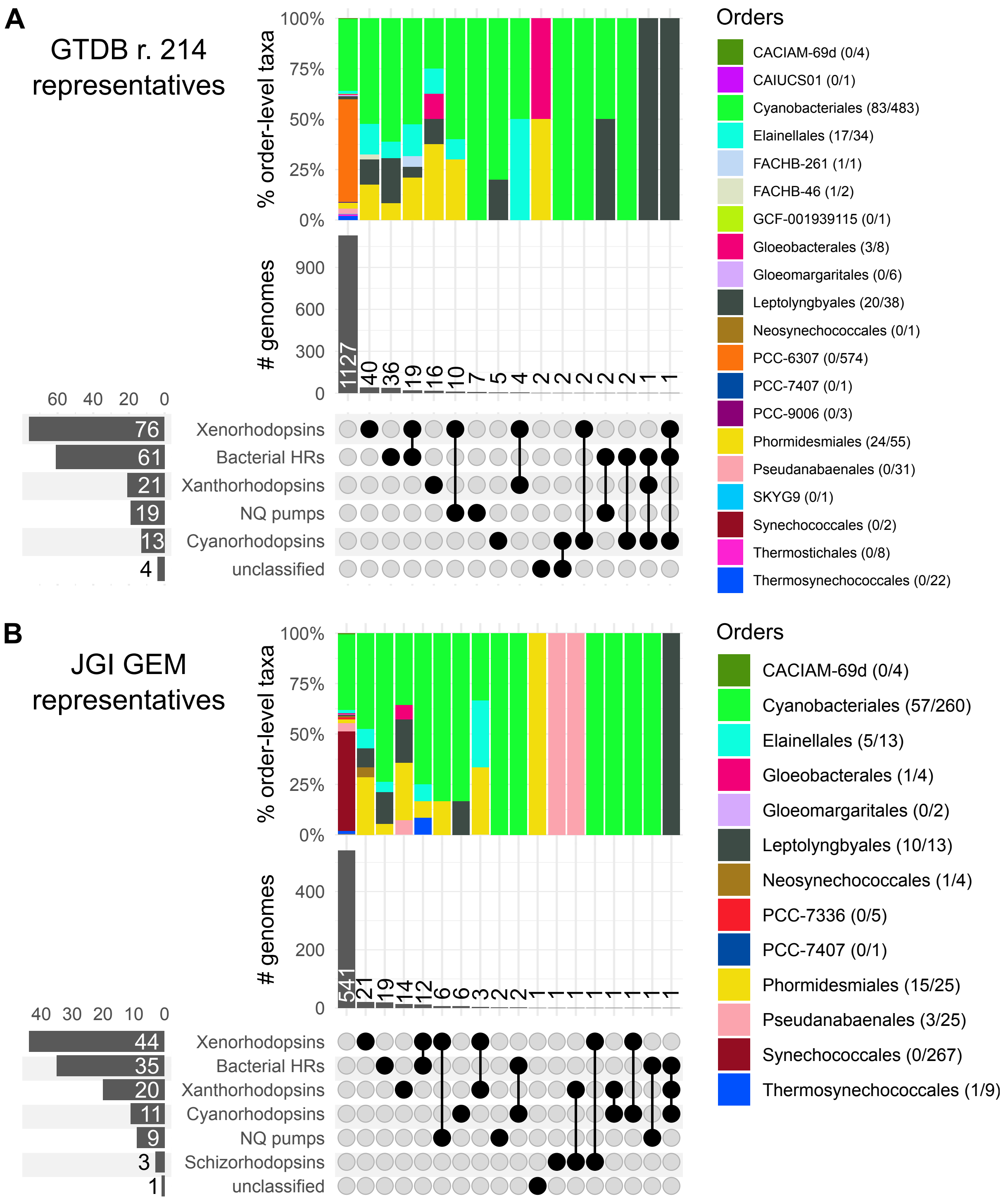


**Supplementary Fig. S1. Distribution of different rhodopsin families in cyanobacteria.** Rhodopsin genes were searched for in representative genomes classified in class "*Cyanobacteriia*" in **A** GTDB r. 214 (species representatives) and **B** JGI GEM (OTU representatives). UpSet plots show the incidence of genomes with no rhodopsin genes (the first column) and genomes coding for rhodopsin genes from a single or multiple rhodopsin families. The genomes are classified to order according to the corresponding database. Counts next to taxa names (orders) indicate numbers of genomes with rhodopsin genes vs. total number of genomes belonging to each taxon.

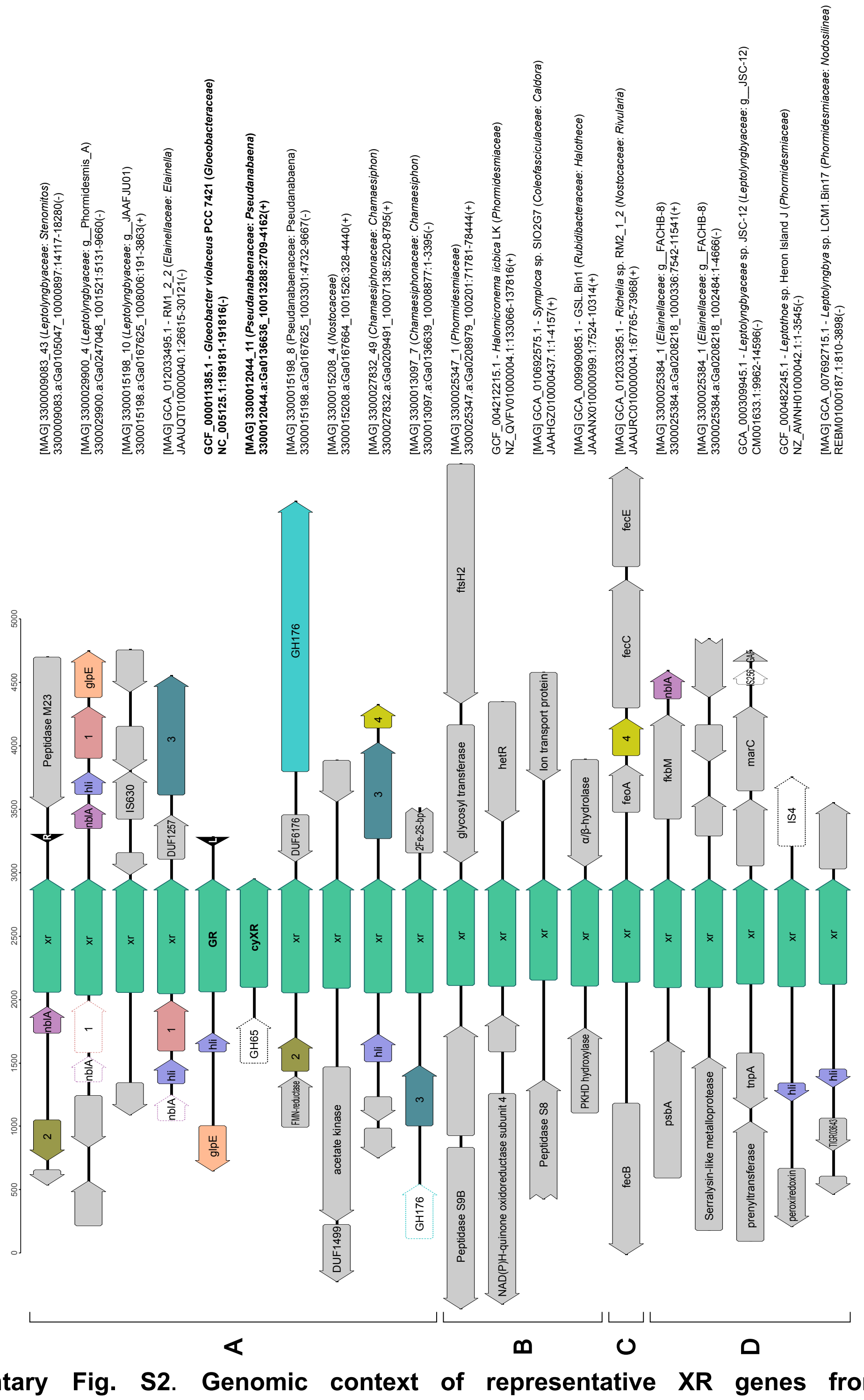


**Supplementary Fig. S2. Genomic context of representative XR genes from diverse cyanobacteria.** ORFs are shown as filled arrows, pseudogenes as white arrows with dashed stroke and tRNA genes as filled black arrowheads. Same color unites homologs with at least two members in the orthogroup with grey fill reserved for singleton genes and black dashed stroke for singleton pseudogenes. Genes are labelled based on predicted function/homology when available. Genes of unknown function but belonging to orthogroups are labelled with numbers. The most frequent genes in the vicinity of the XR genes are *hli* coding for high-light inducible protein and *nblA* coding for proteolysis adaptor NblA. Metadata for each genomic fragment is provided with GTDB taxonomic assignments in parentheses (family and genus if different from the originally assigned genus). The genomic fragments are grouped by XR subclade (see Fig. 1B).

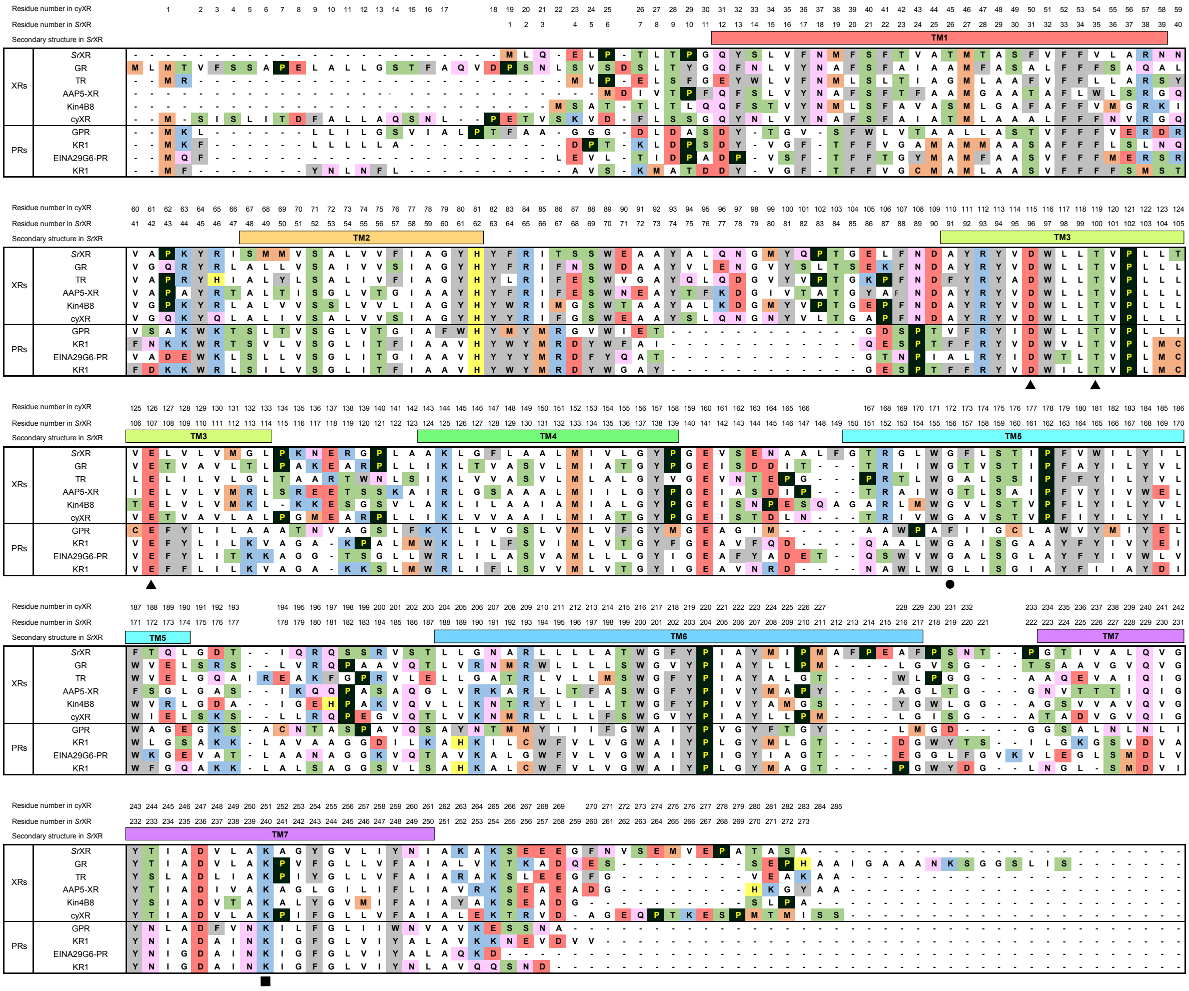


**Supplementary Fig. S3. Multiple amino acid sequence alignments**. The amino acid sequences were aligned using PROMALS3D[1] and MEGA11[2]. *Sr*XR (*S. ruber*), GR (*G. violaceus* str. ATCC 29082), TR (*Thermus thermophilus* JL-18), AAP5-XR (*Sphingomonas glacialis* AAP5), Kin4B8 (uncultured *Bdellovibrionales bacterium*), GPR (*Gammaproteobacterium* EBAC31A08), KR1 (*Dokdonia eikasta* NBRC100814), EINA29G6-PR (uncultured *Flavobacteriales* bacterium), and *Ts*PR (*Tenacibaculum* sp. SG-28) were aligned with the sequence of cyXR from Antarctic cyanobacteria. The residue numbers of *Sr*XR and cyXR are shown on top of the residues, together with the positions of TM helices (indicated by rectangles) based on X-ray crystallographic structure of *Sr*XR (PDB ID: 3DDL)[3]. The three-letter motif, the glycine at the fenestration, and the retinal-binding lysine are indicated by black triangle, circle, and square symbols, respectively.

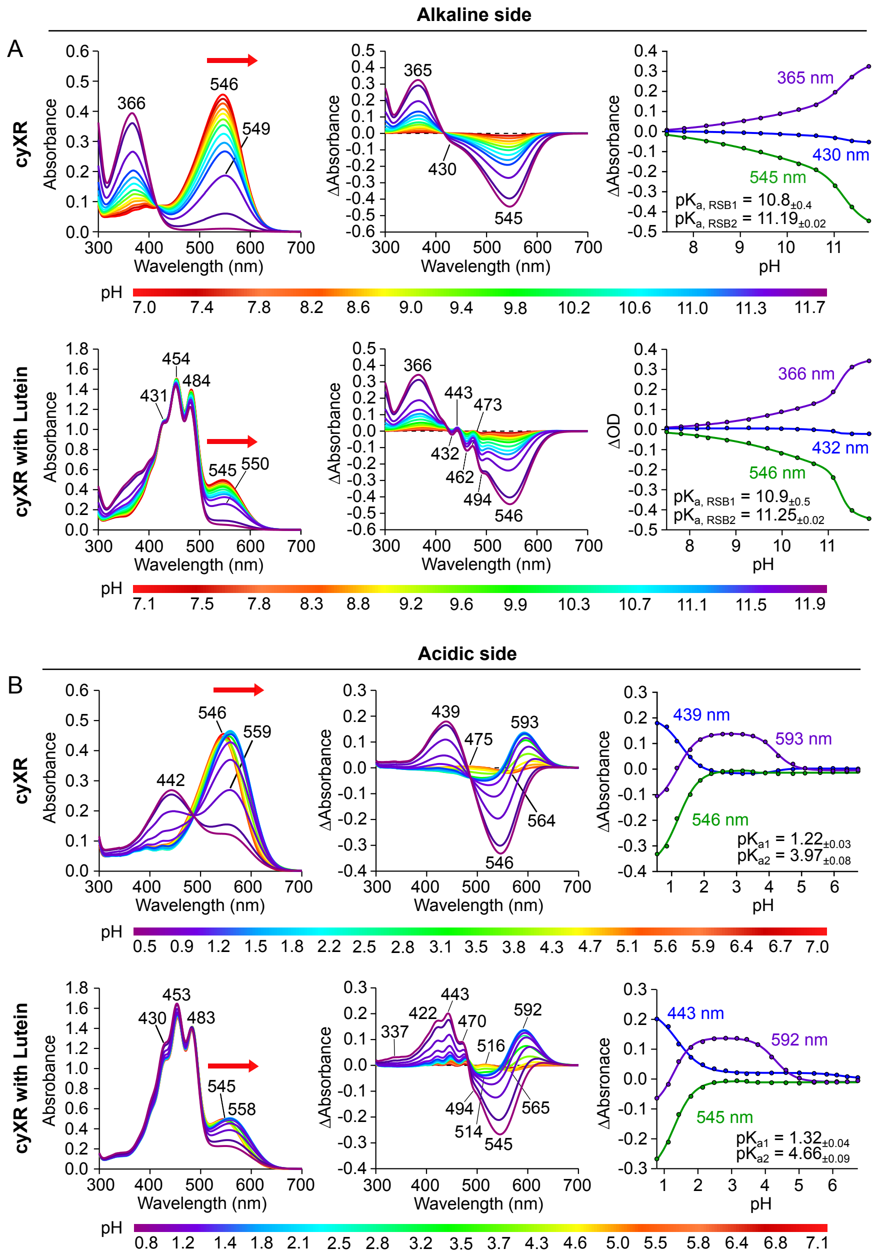


**Supplementary Fig. S4. pH dependence on the absorption**. Absorption spectra (left), different absorption spectra (center), and pH titration curves for the calculation of p$K_a$ (right) were measured depending on **A** acidic and **B** alkaline pH changes. The red-shift in absorption spectra at each pH are indicated by red arrows. The pH titration curves were analyzed using the Henderson–Hasselbalch equation[4].

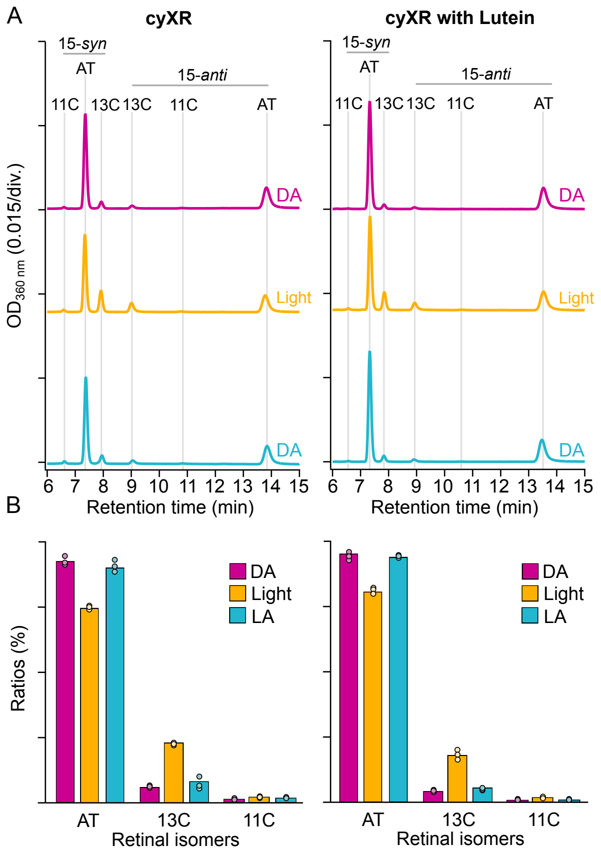


**Supplementary Fig. S5. HPLC analysis of the retinal configuration**. **A** Chromatogram of HPLC analyses and **B** the compositions of the retinal isomers under the dark (DA, violet), light (orange), and light-adapted (LA, blue) conditions, where AT, 13C, 11C, *syn*, and *anti* indicate all–*trans*, 13–*cis*, 11–*cis*, *syn*, and *anti* configurations, respectively. The corresponding data are listed in Supplementary Table S1.

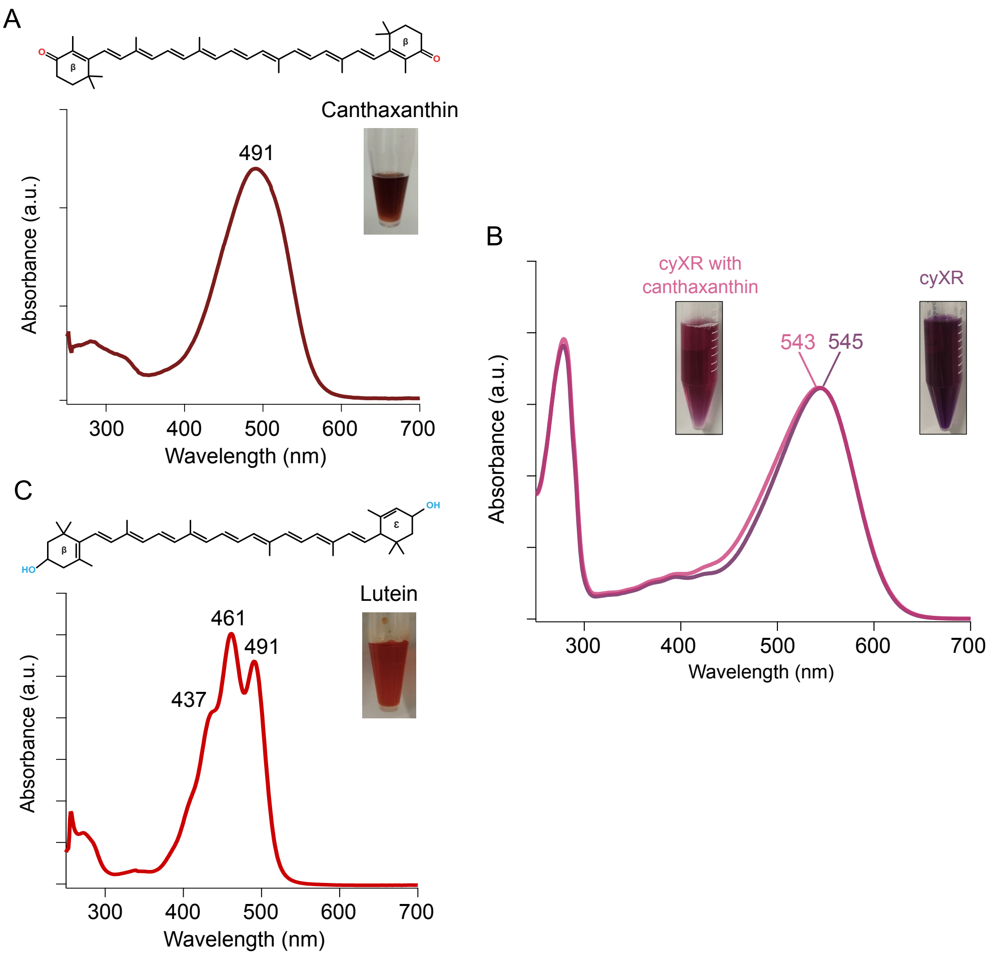


**Supplementary Fig. S6. Absorption spectra of carotenoid antennas**. **A** The chemical structure (top) and UV-vis absorption spectrum (bottom) of canthaxanthin. Canthaxanthin was solved in DMSO; **B** UV-vis absorption spectra of cyXR (violet) and cyXR with canthaxanthin (pink); **C** The chemical structure (top) and UV-vis absorption spectra (bottom) of lutein. Lutein was solved in DMSO. The pictures of the solutions of commercially-available carotenoids and the purified proteins are shown next to the corresponding results.

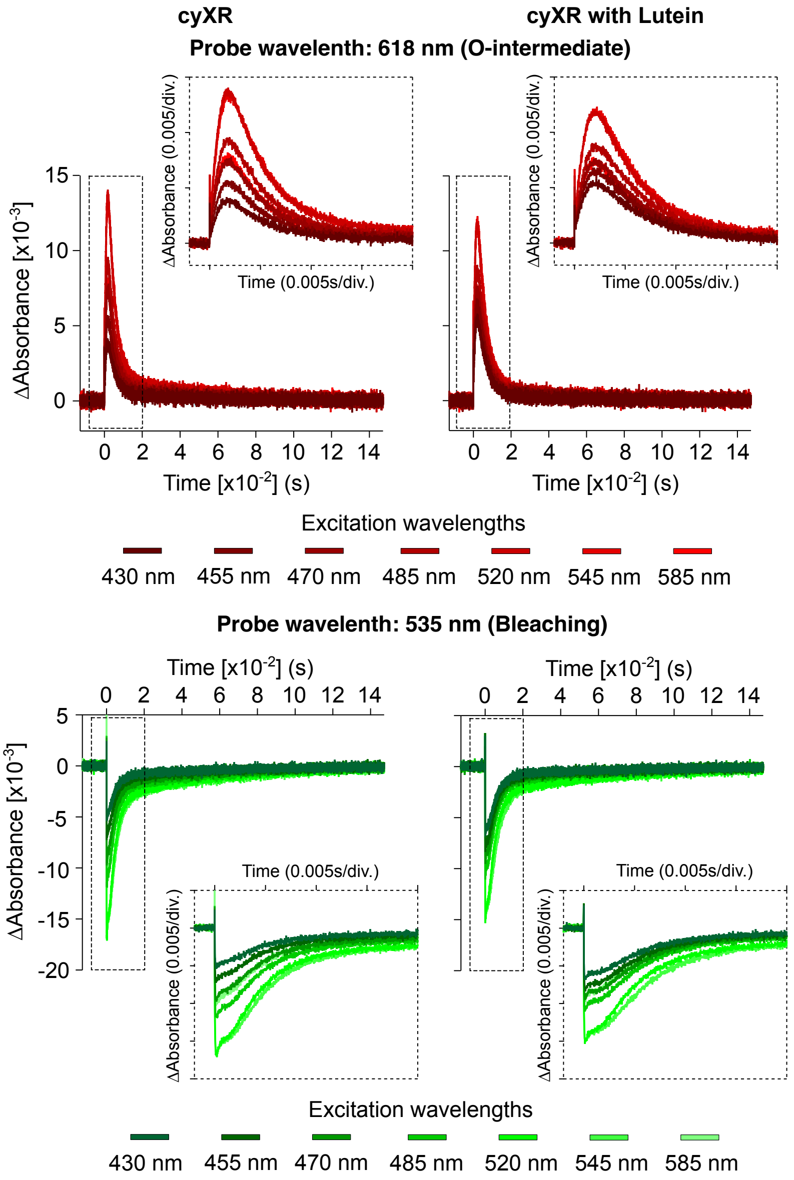


**Supplementary Fig. S7. Excitation energy transfer.** The excitation-wavelength dependence of the transient absorption signals of cyXR without (left) and with (right) lutein probed at 618 (red, top) and 535 nm (green, bottom), representing the O-intermediate accumulation and the initial state bleach, respectively.

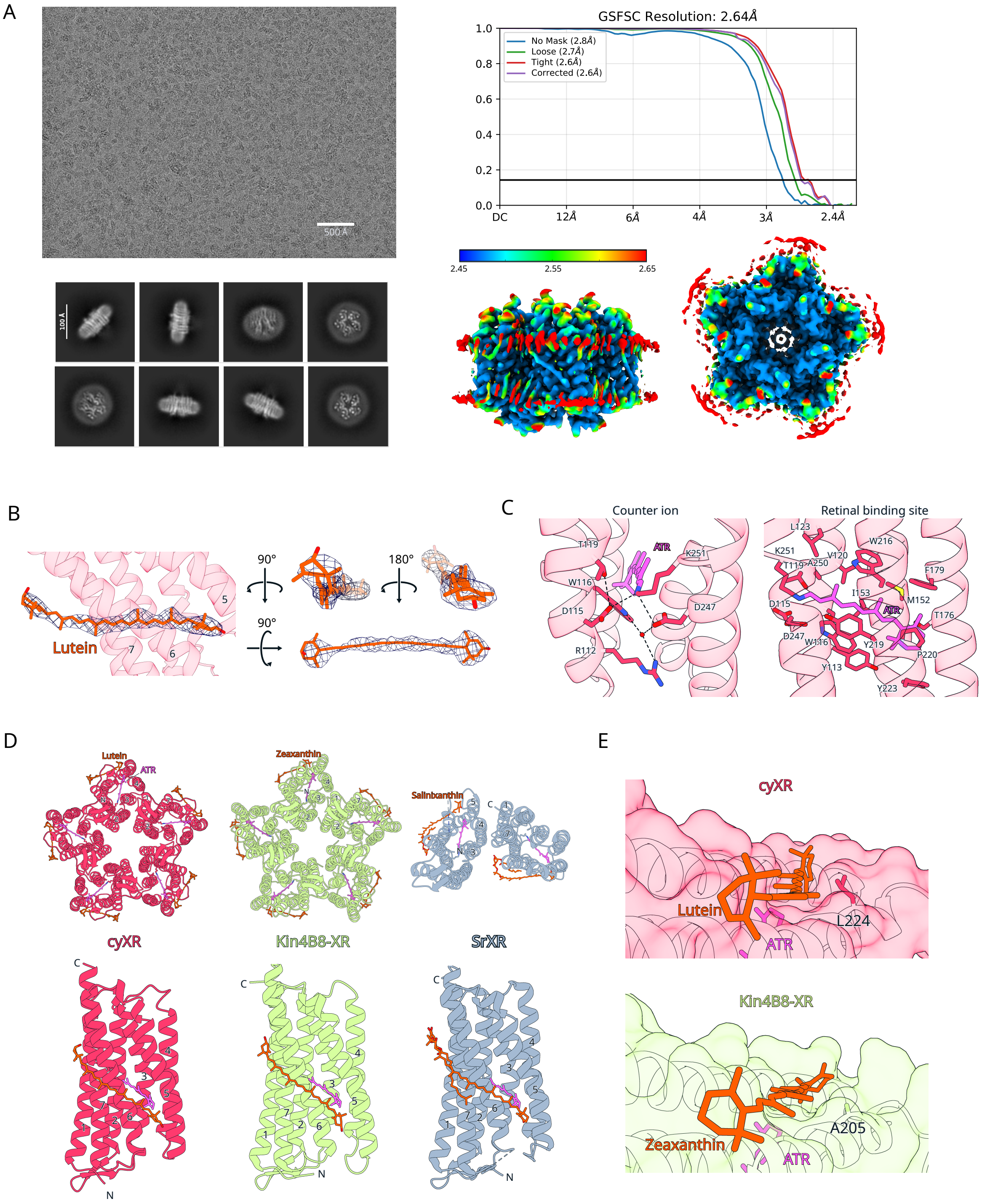


**Supplementary Fig. S8. Structural features of cyXR. A** Cryo-EM single-particle analysis of lutein-bound cyXR; **B** Cryo-EM density of lutein, enabling unambiguous identification of the molecule. Notably, the resolution of the lutein region proximal to the fenestration is sufficient to discern the dimethyl group on its hydroxyl ring; **C** Key residues involved in rhodopsin proton pump motifs in cyXR. Black dashed lines represent hydrogen-bonding interactions, while red spheres denote water molecules. Structural analysis suggests that, unlike in Kin4B8-XR and *Sr*XR, the interaction between D247 and the Schiff base in cyXR is not mediated by a water molecule through hydrogen bonding; **D** Comparison of the oligomeric structures of cyXR, Kin4B8-XR (PDB ID: 8I2Z)[5], and *Sr*XR (PDB ID: 3DDL)[3]. The pentameric structure of cyXR represents its physiological form, in contrast to the previously reported head-to-tail dimer of *Sr*XR; **E** Comparative analysis of the positioning of the carotenoid ring relative to the bulky side chains of L224 in cyXR and A205 in Kin4B8-XR. The steric hindrance imposed by L224 in cyXR causes a shift in the orientation of the lutein ring compared to the zeaxanthin ring in Kin4B8-XR.
